# MEGA-ODE: Learning Biologically Structured and Navigable Continuous Perturbation Dynamics from Sparse Omics

**DOI:** 10.64898/2026.08.05.742921

**Authors:** Yujia Xiang, Yongge Li, Chunyan Tian, Ruichu Gu, Fuchu He, Han Wen, Linhai Xie, Peijie Zhou

## Abstract

Perturbation-omics experiments usually measure only a subset of molecular feature, intervention and time space, leaving many response trajectories, perturbation effects and disease- or differentiation-associated transitions unobserved. Here we present MEGA-ODE, a graph-constrained continuous-time framework for reconstructing sparse dynamic omics landscapes, predicting unmeasured molecular states and prioritizing virtual perturbations toward defined biological endpoints. MEGA-ODE integrates molecular-network priors, graph neural ordinary differential equations and context-adaptive mixture-of-experts routing. In L1000 transcriptomic perturbations and CPPA proteomic drug-response data, MEGA-ODE improved held-out-feature and unseen-perturbation prediction over baseline methods, and in SARS-CoV-2 infection time-series data it remained competitive for future-time-point forecasting. In a COVID-19 patient cohort, predicted intermediate profiles improved retrospective disease-stage stratification relative to observed profiles alone, while expert programs highlighted immune and inflammatory signals associated with severity. Across the MAPK drug-response and stem-cell differentiation case studies, graph- and expert-level attributions prioritized perturbation-associated MAPK edges, developmental regulators and TF-target relationships supported by independent promoter-proximal ChIP-seq overlap. In hESC-to-definitive-endoderm differentiation, MEGA-ODE prioritized candidate transcription-factor perturbations predicted to shift 12-36 h profiles toward 96 h definitive-endoderm marker signatures, framing trajectory navigation as a concrete hypothesis-generation task. Together, these results support biologically structured continuous-time modeling for prediction, interpretation and virtual-perturbation prioritization from sparse temporal omics data.

## Introduction

Dynamic biological processes are rarely observed and measured at the temporal or perturbational resolution where they unfold. In drug response, disease progression and cell differentiation, molecular states change continuously, whereas high-throughput measurements typically capture a limited set of genes or proteins, a small number of perturbations and a sparse series of time points. This mismatch leaves many biologically important states unmeasured, including responses of unprofiled molecules, effects of untested perturbations and transient states between sampled time points. As a result, temporally-resolved omics datasets require models that can extrapolate beyond the measured experiment while preserving links to prior biological knowledge and producing hypotheses that can be followed experimentally.

Dynamic omics studies commonly face three forms of missingness. First, measurements often cover only part of the relevant molecular state space: genes or proteins with potential roles in the response may be absent from the assay, although their trajectories can be important for interpreting pathway activity and regulatory programs^1,3,4^. Second, experimental designs can include only a limited subset of perturbations, drugs and combinatorial conditions^3,5^. Third, biologically meaningful transitions may occur between sampled time points, leaving intermediate states unobserved^1,6^. Together, these limitations make it difficult to infer how molecular systems evolve outside the measured feature, perturbation and time axes, raising a central methodological question: how can unmeasured components of dynamic biological responses be estimated from sparse and mosaic experimental observations?

Existing methods provide partial solutions to this problem, yet most of them focus on extrapolation along a single dimension. Perturbation-prediction methods can estimate the state induced by a drug or genetic intervention, but they often focus on state mapping and do not represent the continuous transition through intermediate molecular states^7,8^. Dynamic models, including latent ODEs and patient-state attention models, can handle time-series data, but many learn trajectories without an explicit connection to signaling, regulatory or physical interaction networks^6,9,10,11,12^. Graph-based models can use molecular priors, for example protein interactions or gene functional networks, and are seldom evaluated in a framework that combines graph structure, time extrapolation and cross-perturbation generalization^5,13,14^. At the model level, deep message passing can blur node-specific signals, and conventional Neural ODE parameterizations may be too restrictive for heterogeneous biological dynamics^15,16^. These limitations motivate a continuous model that uses molecular networks to guide prediction across unmeasured molecules, unseen perturbations and unsampled time points.

To address this task, we introduce MEGA-ODE (Mixture-Enhanced Graph Attention ODE), a continuous-time framework that combines molecular-network priors, mixture-of-experts dynamics and graph attention. Its design follows three biological considerations. First, sparse omics observations often permit trajectories that explain the data equally well yet violate biological plausibility. MEGA-ODE therefore uses curated molecular-network priors to constrain candidate transitions learnt by the Neural ODE to reported regulatory, physical or functional relationships^13,14^. Second, biologically plausible responses can arise through multiple dynamical programs whose contributions vary across conditions and time. Drug exposure, infection and differentiation can combine stress signaling, feedback, lineage programs and recovery, so MEGA-ODE represents each response as a mixture of graph ODE experts with context-dependent gating, to overcome the limited expressive capacity of a single ODE parameterization^17,18^. Third, the use of multiple graph ODE experts increases the expressive capacity of the model but also makes the learned dynamics more difficult to interpret. Because biologists seek biological insight in addition to predictive accuracy, MEGA-ODE employs graph attention to highlight influential molecular interactions and expert usage patterns, providing molecular clues for experimentally testable hypotheses^19^.

Accordingly, we evaluated MEGA-ODE at three complementary levels. First, we asked whether the model could reconstruct unmeasured biological states from sparse experimental observations by assessing its ability to generalize across three forms of missing information. Feature-masking experiments tested whether network-constrained dynamics could infer responses for molecules withheld from training, held-out drug experiments tested generalization to perturbations absent from training, and future-timepoint experiments tested whether continuous dynamics could support prediction across unsampled time windows. Second, we asked whether the well-performing models could also provide biological insight into their predictions. This layer of evaluation examined pathway behavior in perturbation datasets, severity-associated programs in patient time series and regulator programs during stem-cell differentiation. Third, we asked whether the learned dynamics could guide biological trajectories toward desired endpoints by prioritizing virtual transcription-factor perturbations during stem-cell differentiation. Across these settings, MEGA-ODE improved masked-feature and held-out perturbation prediction, remained competitive in future-timepoint forecasting, and prioritized candidate perturbations predicted to steer differentiation toward desired endpoints. Its attributions highlighted MAPK-related associations, disease-severity-associated programs, differentiation-associated regulators and selected TF-target links with independent ChIP-seq promoter-overlap support. Together, these results support biologically structured continuous-time modeling for prediction, interpretation and virtual-perturbation prioritization from sparse dynamic omics data.

## Results

### MEGA-ODE: a biologically structured dynamics model with modular expert programs

MEGA-ODE takes two inputs: a prior molecular interaction graph and a time-ordered omics matrix whose rows represent samples, pseudo-bulk states or time-point states and whose columns represent measured genes or proteins. To address the challenge of modeling dynamic responses under diverse perturbations, MEGA-ODE integrates these inputs through structured biological priors and continuous-time dynamics. The model comprises three components: (i) a graph encoder that propagates measured omics features over the supplied biological network, such as protein-protein or gene-gene associations; (ii) a mixture-of-experts (MoE) framework that decomposes heterogeneous dynamics into specialized modules, with a gating network that weights expert contributions for each context; and (iii) a graph-based neural ODE backbone that models continuous expression evolution (**Fig. 1A, Supplementary Fig. 1**).

**Fig. 1.**
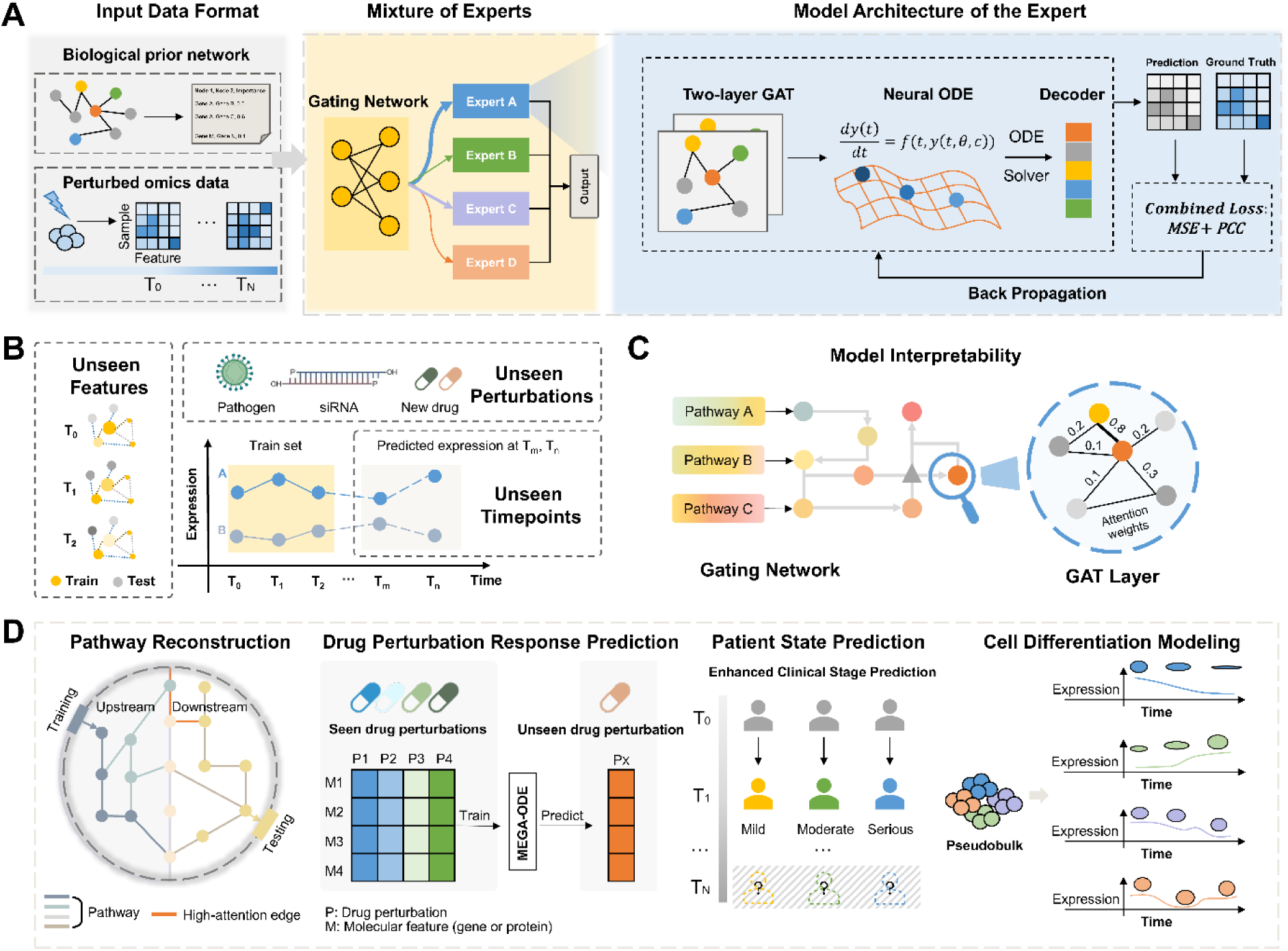
Model framework of MEGA-ODE. (A) Model architecture of MEGA-ODE. MEGA-ODE takes two inputs: (i) a prior molecular interaction graph, such as protein-protein interaction (PPI) networks from STRING or gene-gene interaction networks from HumanNet, and (ii) a time-ordered omics matrix under perturbation. In the matrix, rows denote biological samples, patient samples or pseudo-bulk state/time-point profiles, and columns denote measured genes or proteins. The model comprises a mixture-of-experts framework with a gating network and a graph attention network (GAT)-based neural ordinary differential equation module, allowing for the integration of biological priors with dynamical system modeling. (B) Generalization tasks evaluated in this study. MEGA-ODE is assessed under three held-out settings: prediction on unseen features, unseen perturbations, and unseen timepoints. (C) Graph- and expert-level interpretability of MEGA-ODE. (D) Application scenarios of MEGA-ODE.

The design follows a staged rationale. The prior graph biases the learned dynamics toward reported molecular associations, improving biological plausibility relative to unconstrained trajectory fitting. Because a fixed graph can also restrict expressiveness when multiple processes are superimposed, the MoE layer provides parallel dynamical programs that can be combined in a context-dependent manner. The neural ODE backbone then models expression change as a continuous-time flow between observed samples, and graph attention supplies edge-level quantities that can be interpreted post hoc. MEGA-ODE predicts future profiles and returns expert weights and graph-attention patterns that can be compared with pathway, clinical and chromatin-binding evidence. The model is trained end-to-end by minimizing losses that penalize discrepancies between predicted and observed molecular states across sampled time points.

A core objective of MEGA-ODE is to generalize beyond the experimental conditions seen during training. To this end, we designed a multi-level evaluation pipeline encompassing three types of held-out scenarios: (i) unseen features, to predict the trajectories of genes or proteins that were masked during training; (ii) unseen perturbation, to predict system response to drugs or other external stimuli not included in the training set; and (iii) unseen time points, to forecast molecular states at future or intermediate time points lacking direct measurements (**Fig. 1B**). Across these evaluation axes, the gating network provides a mechanism for combining expert outputs; subsequent benchmarks test whether this design is associated with improved held-out prediction. For example, perturbations with related downstream responses may reuse similar expert combinations, supporting a model-based route to held-out perturbation prediction.

Beyond predictive accuracy, we investigated whether learned representations align with known biological annotations (**Fig. 1C**). The interpretability of the model arises from how it organizes transcriptional dynamics during multi-timepoint prediction. The model represents temporal changes through multiple expert-weighted components. The gating network assigns soft weights to expert modules, and these modules can be associated post hoc with enriched pathways and stage- or condition-associated programs. Within each expert, candidate relationships are further summarized through a graph attention mechanism over the supplied gene-gene or protein-protein network. Attention weights provide model-derived prioritization of genes and edges within a given context, highlighting candidate associations for follow-up analysis. Because attention is learned separately within each expert, the same gene can carry different model importance across expert programs, consistent with the context-dependent roles of genes in biological systems. Together, this hierarchical structure enables the model to represent differentiation-associated expression dynamics as a composition of expert-weighted programs, separating coarse program selection from finer-grained graph-based edge prioritization.

Consequently, MEGA-ODE is applicable across diverse biological contexts (**Fig. 1D**). Inferring trajectories of unobserved genes or proteins enables graph-constrained propagation of molecular responses along interaction networks. By capturing system responses to novel drugs or stimuli, it supports held-out pharmacological prediction in settings where related perturbations share response structure. By modeling continuous temporal dynamics, it forecasts molecular states and patient trajectories. Beyond prediction, the mixture-of-experts architecture enables learned components to be annotated post hoc with coherent signals such as differentiation-associated, apoptosis-associated, or cell-cycle-associated programs.

### MEGA-ODE achieves robust prediction of masked molecular features

To evaluate prediction of unobserved features (**Fig. 2A**), we benchmarked MEGA-ODE on three perturbation datasets: transcriptomics under biological agents, transcriptomics under drug treatment, and proteomics under compound perturbation. In the reported splits, MEGA-ODE achieved the highest or near-highest Pearson correlation and lower MAE than the tested recurrent and ODE-based baselines, including models with or without graph priors and expert modularity (**Fig. 2B**, **Supplementary Fig. 2**). NODE-based models performed competitively, but combining graph encoding with the mixture-of-experts architecture yielded improved accuracy, suggesting that the model benefits from both graph-informed structure and modular decomposition in these tasks.

**Fig. 2.**
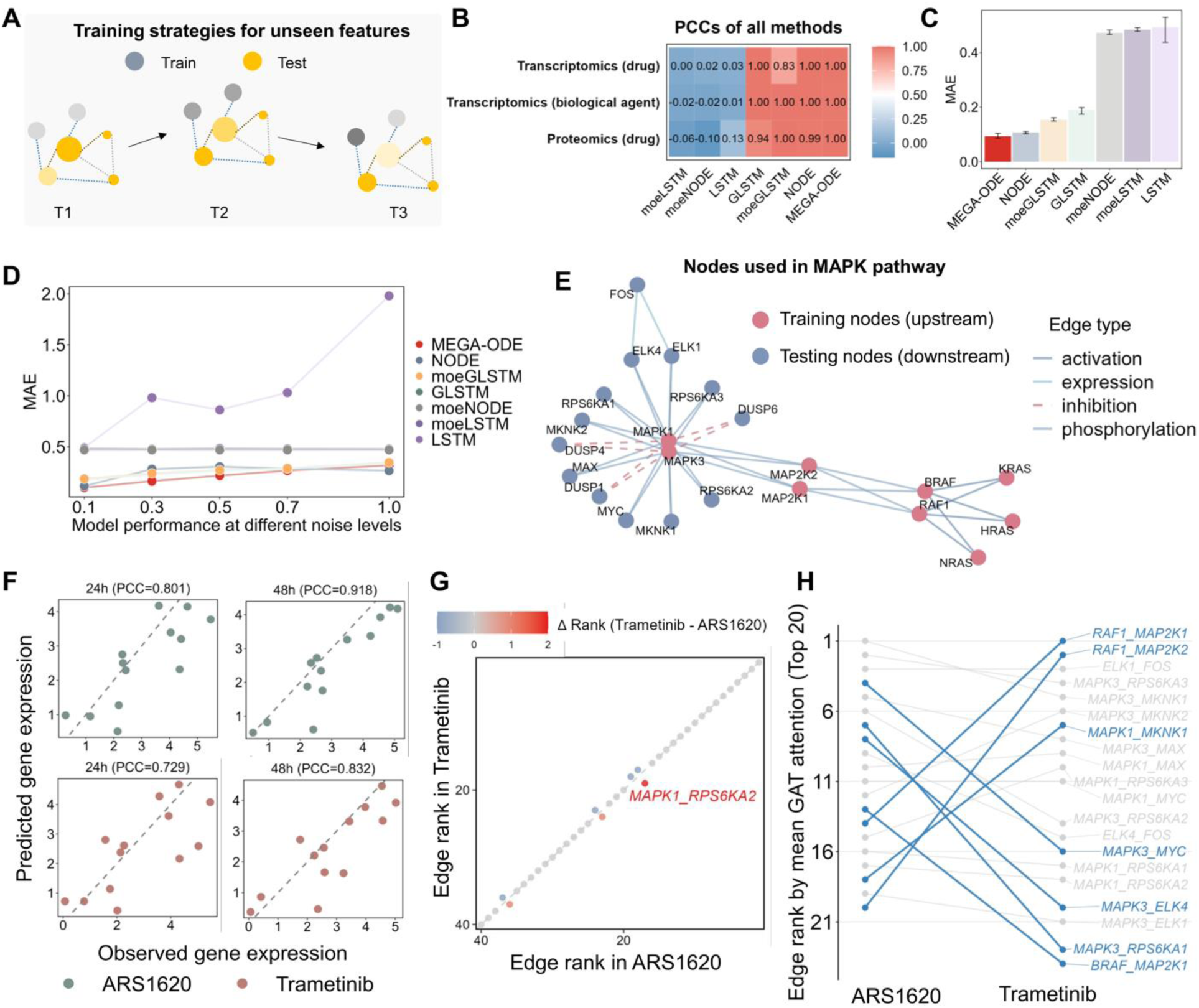
Generalization performance, robustness, and interpretability of MEGA-ODE under unseen-feature settings. (A) Schematic of the unseen-feature prediction task. (B) Comparative evaluation across perturbational omics datasets. Model performance is evaluated across three perturbational datasets, including L1000 transcriptomic profiles under biological agent perturbations, drug-perturbed transcriptomes of the MCF cell line, and the cancer perturbed proteomics atlas (CPPA). A fixed masking ratio of 0.5 is applied, and performance is assessed using the Pearson correlation coefficient. (C) Prediction error on drug-perturbed proteomics data. MAE is evaluated on the CPPA dataset under the unseen-feature setting with a masking ratio of 0.5. (D) Performance robustness under additive noise. Model performance is assessed by progressively adding Gaussian noise to the input data. The x-axis denotes the standard deviation of the added noise, and the y-axis reports MAE. (E) MAPK pathway gene set used for biological analyses. MAPK1 (ERK2) and MAPK3 (ERK1), together with their upstream regulators, are treated as training nodes, while post-ERK downstream genes are masked as testing targets. (F) Prediction performance on masked downstream nodes. MEGA-ODE is trained on 4-h expression profiles after ARS-1620 or Trametinib treatment and predicts downstream gene expression at 24 and 48 h. (G) Cross-perturbation comparison of edge-attention ranks among upstream training nodes. Each point represents the rank of the same edge in the ARS-1620 and Trametinib datasets. (H) Cross-perturbation comparison of downstream testing-node edge-attention ranks, highlighting post-ERK edges whose relative importance differs between ARS-1620 and Trametinib.

To assess numerical accuracy, we computed the MAE on held-out proteomic features. MEGA-ODE showed lower MAE than the tested recurrent and ODE-based baselines in this held-out proteomic setting (**Fig. 2C**). We next evaluated robustness to partial graph knowledge by masking increasing proportions of edges in the prior network. While all graph-based models degraded with edge removal, MEGA-ODE maintained comparatively lower error and flatter degradation curves, suggesting stability against prior incompleteness in this analysis (**Supplementary Fig. 3**). We further tested resilience to feature noise by adding Gaussian perturbations. MEGA-ODE showed relatively small changes in MAE across the tested noise levels (**Fig. 2D**). These results support MEGA-ODE’s use for predicting unobserved features under the evaluated structural- and measurement-noise settings.

### MEGA-ODE prioritizes pathway-level signaling-consistent structure

We next examined whether model-derived edge rankings captured expected structure in a well-characterized MAPK signaling task. We analyzed time-resolved transcriptomes from NCI-H358 cells treated with either the covalent KRAS(G12C) inhibitor ARS-1620 or the MEK inhibitor Trametinib^20^. Because both drugs perturb the KRAS-RAF-MEK-ERK axis at different positions, we split the pathway around ERK, using MAPK1 (ERK2), MAPK3 (ERK1), and their upstream regulators as observed training nodes, whereas genes downstream of ERK were masked during training and served as held-out test nodes for model validation (**Fig. 2E**)^21,22^.

MEGA-ODE predicted masked downstream gene expression at 24 and 48 h after training on the 4-h response, both for ARS-1620 and Trametinib perturbations (**Fig. 2F**). This design tests whether early upstream pathway information can be propagated through the known graph to predict later post-ERK transcriptional outputs. The results are consistent with graph-mediated transfer from the shared KRAS-RAF-MEK-ERK trunk to downstream target genes, despite those downstream genes being withheld from the training loss.

We next compared graph-attention ranks between the ARS-1620 and Trametinib models. Among upstream training nodes, edge rankings were highly concordant across the two perturbations (**Fig. 2G**). This is consistent with the expected biology: although ARS-1620 acts at KRAS and Trametinib acts at MEK, both compounds converge on the same upstream MAPK signaling trunk before ERK activation. One edge, MAPK1-RPS6KA2, ranked slightly higher in ARS-1620 than in Trametinib. Because the ranks are defined within each independently trained model, the modest MAPK1-RPS6KA2 shift provides a qualitative indication of perturbation-associated edge reweighting.

In contrast, downstream testing-node edge ranks differed more substantially between the two perturbations (**Fig. 2H**). These differences indicate that the two inhibitors are associated with distinct post-ERK transcriptional patterns despite sharing a common upstream axis. The ARS-1620 model assigned high importance to the MAPK3-MYC edge, matching the original Cell study’s report of a prominent MYC-associated expression cluster after KRAS(G12C) inhibition^20^. The model-prioritized edges are provided in **Supplementary Table 1**. This comparison evaluates concordance between selected model-prioritized edges and perturbation-linked transcriptional evidence at the tested time points. Together, these analyses show that MEGA-ODE recovered shared upstream MAPK structure and perturbation-associated downstream edge rankings in this pathway task.

Together, these results support the conclusion that MEGA-ODE can impute masked molecular features and prioritize signaling-consistent pathway associations using graph-informed dynamics and modular decomposition.

### MEGA-ODE generalizes to unseen drug perturbations and reveals expert-associated programs

Generalizing predictive models to novel perturbations is critical for enabling prospective inference in systems pharmacology. To assess how MEGA-ODE performs under unseen perturbations, we evaluated its ability to predict molecular responses to previously unseen drugs using proteomic profiles from the Cancer Perturbed Proteomics Atlas (CPPA)^4^ (**Fig. 3A**). We selected structurally diverse compounds with varying mechanisms of action (MOAs) for leave-one-drug-out cross-validation. Drug similarity was quantified using Tanimoto coefficients based on chemical structure, revealing a broad spectrum of pairwise similarities (**Fig. 3B**, Methods).

**Fig. 3.**
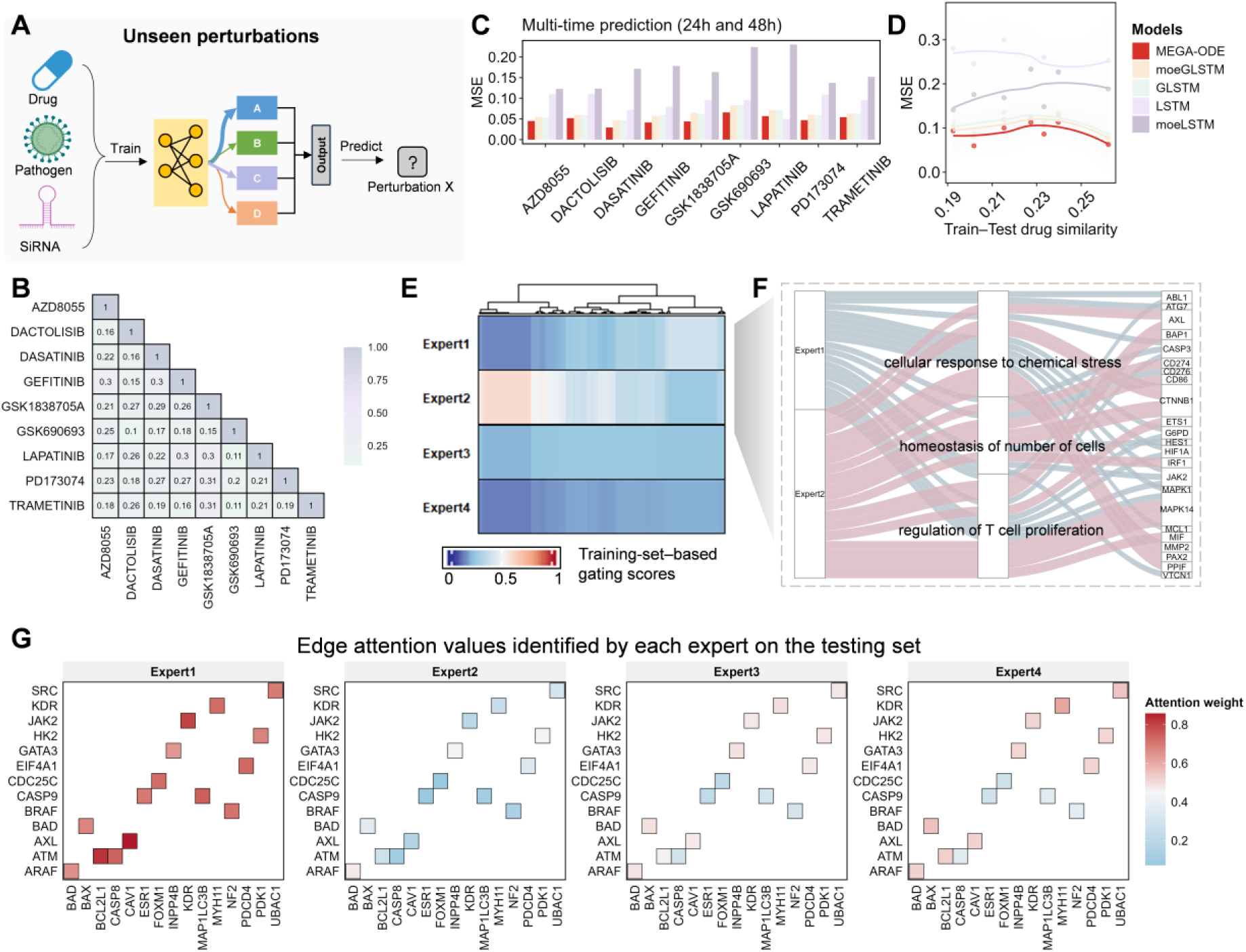
Generalization to unseen drug perturbations and expert-level interpretation from MEGA-ODE. (A) Schematic of the unseen-perturbation prediction task. Models are trained on a subset of drug perturbations and evaluated on a held-out drug that is not observed during training. (B) Chemical similarity among the nine drugs used in the drug-masking analysis, quantified using the Tanimoto coefficient. (C) Prediction performance on unseen drug perturbations. A leave-one-drug-out evaluation is performed on a total of nine drugs, where models are trained on eight drugs and evaluated on the remaining unseen drug. (D) Model performance and train-test drug similarity. The x-axis represents the mean chemical similarity between the held-out test drug and the drugs used for training, while the y-axis reports the mean squared error (MSE) of model predictions. (E) Protein-level gating weight distributions learned from training data. Columns correspond to proteins, rows correspond to the four experts, and values indicate the gating scores assigned to individual proteins. (F) Gene Ontology enrichment analysis of expert-specific gene sets. (G) Expert-specific interaction patterns under an unseen drug perturbation. For the unseen drug Trametinib, graph attention weights are extracted from the trained model on the testing set. Important edges identified by the attention mechanism are shown for each expert, reflecting expert-specific interaction patterns under previously unseen perturbations.

Across most of the held-out drug conditions, MEGA-ODE achieved lower MSE in predicting future protein expression than the tested recurrent baselines with or without graph priors and expert decomposition (**Fig. 3C**). Nine held-out drugs are shown in the main panel to provide a complete view of model performance across the evaluation set. Within this nine-drug CPPA leave-one-drug-out analysis, MEGA-ODE maintained comparatively low error across the observed range of train-test structural similarities (**Fig. 3D**). These results indicate that MEGA-ODE’s mixture-of-experts structure can support held-out drug prediction in this dataset without relying solely on molecular similarity.

To clarify the pharmacological diversity of the modeling task, we categorized the drugs by their known MOAs (tyrosine kinase signaling, including receptor tyrosine kinase programs such as VEGFR signaling in cancer^23^; metabolic regulation; PI3K-AKT signaling; and cell-cycle control; **Supplementary Fig. 4**). We further evaluated a cross-MOA setting in which two MOA groups were used for training and the remaining MOA group was held out for testing across six train-validation-test arrangements. After grouping runs by the held-out MOA, MEGA-ODE showed higher PCC than the tested baselines in these CPPA cross-MOA splits, supporting limited evidence for held-out MOA prediction within this dataset (**Supplementary Fig. 5**).

We then examined whether MEGA-ODE captures this functional heterogeneity through expert specialization. The gating score matrix in **Fig. 3E** is protein-level: each column corresponds to one protein, and each row reflects the weight assigned to one expert for that protein. The observed soft composition indicates that MEGA-ODE combines latent dynamics across experts while allowing individual proteins to preferentially engage specific expert programs. Such flexibility may support generalization to unseen drug perturbations.

Despite this diffuse routing pattern, distinct functional signatures emerged across experts. Aggregating gating weights across the top-weighted gene sets of each expert and mapping them to Gene Ontology categories showed enrichment for biologically coherent pathways. Expert 1 predominantly weighted proteins involved in chemical stress responses, while Expert 2 was enriched for targets associated with T-cell regulation and the homeostasis of cell numbers (**Fig. 3F**). These patterns are consistent with expert modules being annotatable by biological functions.

This functional annotation was further supported by graph-attention patterns within each expert module. For example, Expert 1 highlighted the ATM-BCL2L1 interaction (**Fig. 3G, Supplementary Fig. 6**), linking a DNA-damage and cellular-stress kinase to an anti-apoptotic BCL-2 family regulator; this pair is consistent with Expert 1’s enrichment for chemical-stress programs and apoptotic effectors. Expert 2 and Expert 3 emphasized immune- and cytokine-signaling-associated nodes, including FOXP1, GATA3 and JAK2; JAK2 is a core component of JAK-STAT signaling and has been linked to tumor-growth programs in breast cancer cells^24^. Expert 4 highlighted RAS-ERK components, including BRAF and MAPK1 (**Fig. 3G, Supplementary Fig. 6**), consistent with established roles of RAS-ERK signaling in cancer and cell behavior^21,22^. Together, these attention patterns support the interpretation that held-out drug predictions are accompanied by biologically annotated expert programs.

Together, these results support the conclusion that MEGA-ODE can predict held-out drug responses across diverse CPPA perturbations while internally organizing predictions into expert-associated programs and pathway-associated attention patterns.

### Temporal forecasting is associated with improved disease-state classification after data augmentation

Many translational applications require modeling molecular changes over time, particularly when dynamic trajectories reflect infection or disease progression. We first evaluated whether MEGA-ODE can extrapolate across unobserved time points in SARS-CoV-2 infection time-series data, and then asked whether model-generated profiles could support disease-state classification in a longitudinal COVID-19 patient cohort (**Fig. 4A**).

**Fig. 4.**
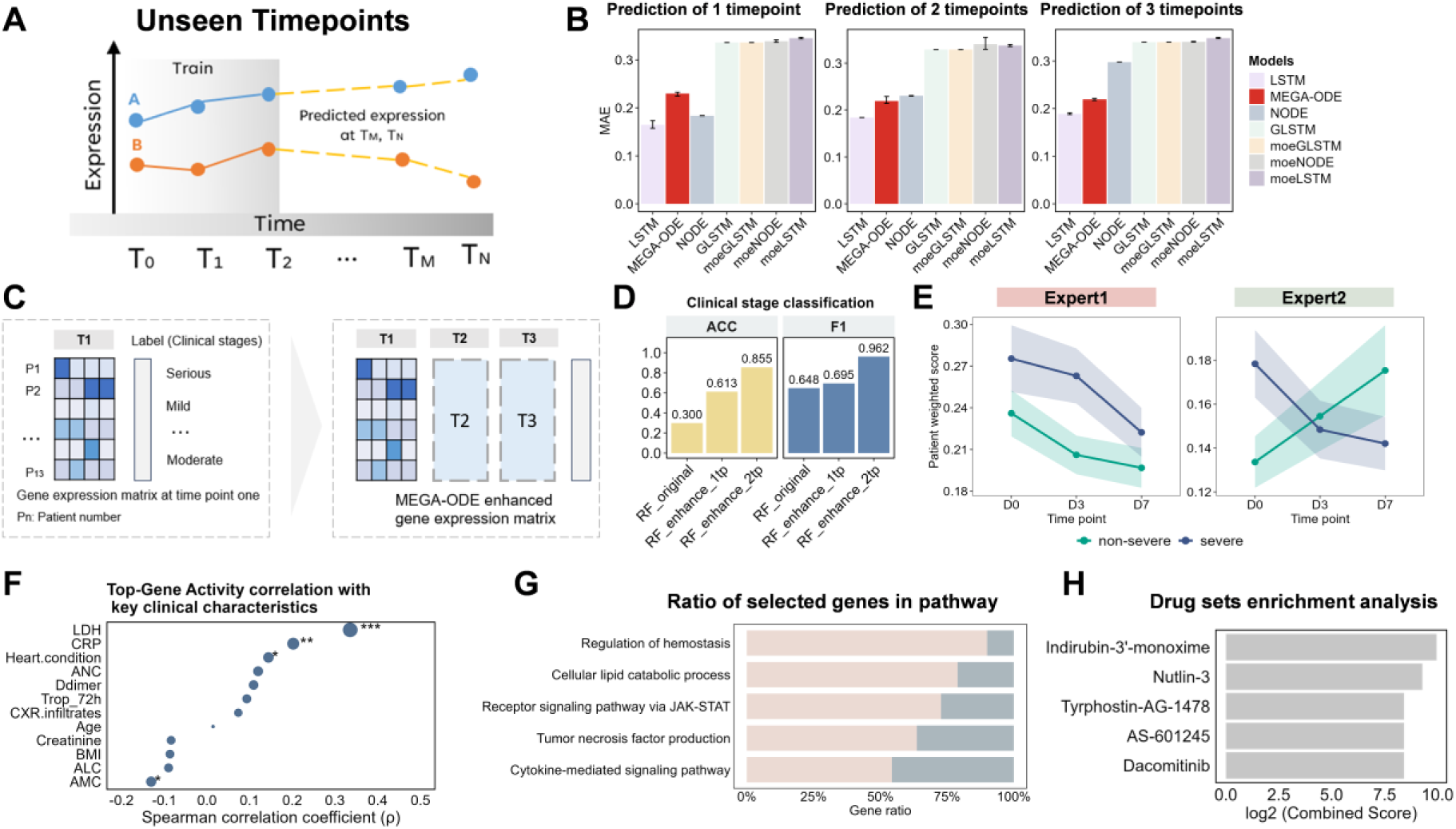
Unseen timepoint prediction. (A) Schematic of the unseen-timepoint prediction task. (B) Model performance when predicting different numbers of future time points. (C) Example of disease-state classification after data augmentation generated by MEGA-ODE. (D) Classification performance of a random forest predictor before and after MEGA-ODE-based data augmentation. Accuracy (ACC) and F1 score are reported for disease-state prediction using the mild, moderate and severe/serious labels in the source cohort. Bar colors distinguish the original-data classifier from classifiers trained after augmentation with two or three MEGA-ODE-predicted future time points, and grouped bars report ACC and F1 under the same patient-level split. (E) Temporal profiles of Top-Gene Activity (TGA) derived from gating scores. Lines indicate mean TGA across patients, shaded areas represent the range across individuals, and colors distinguish disease-severity groups. (F) Spearman correlation between TGA and clinical phenotypes. (G) Gene set enrichment analysis based on attention-ranked genes. Stacked bar colors indicate the two expert-associated selected-gene sets contributing to each pathway gene ratio. (H) Drug-signature enrichment analysis based on the top-ranked gene sets by node importance.

The benchmark dataset contains measurements collected at 0, 1, 2, 3, 6 and 12 h. Using partial observations (one to three time points) from SARS-CoV-2-infected human lung epithelial cells^25^, we compared seven models in predicting future gene expression. MEGA-ODE showed competitive performance across forecasting horizons, with prediction errors comparable to the best-performing baseline when extrapolating two or three time points (**Fig. 4B**). LSTM slightly outperformed MEGA-ODE in the short-horizon setting, so we interpret this benchmark as evidence of competitive temporal forecasting. The advantage of MEGA-ODE in this setting is that its forecasts are coupled to graph attention and expert-level programs that can be interrogated biologically. Under a controlled setting where the training set was fixed and increasing levels of noise were added to the test data, all models exhibited performance degradation; however, the MAE gap between MEGA-ODE and the best-performing LSTM progressively narrowed, with the two curves nearly overlapping at higher noise levels (**Supplementary Fig. 7**).

We next examined whether MEGA-ODE-generated profiles could improve disease-state prediction in the COVID-19 patient cohort^26^. We augmented observed expression matrices with MEGA-ODE-predicted future time points and trained a random forest classifier to infer disease stage (**Fig. 4C**). In this small cohort analysis, incorporating MEGA-ODE-predicted future time points increased cross-validated accuracy and F1 score relative to observed data alone (**Fig. 4D**), indicating that the predicted features contain information useful for disease-state classification in this cohort and motivating validation in independent longitudinal cohorts.

### Expert-specialized gene prioritization reveals molecular correlates of disease severity

Beyond forecasting, a key goal in time-series modeling is to identify interpretable molecular programs that vary across patient subpopulations. To probe the latent structure learned by MEGA-ODE, we analyzed expert-specific gene weighting in a COVID-19 patient cohort sampled over three timepoints (days 0, 3, and 7)^27^. This cohort contains more samples and richer clinical phenotypes than the benchmark dataset, enabling links between model-derived gene programs and disease symptoms.

Two expert modules emerged as dominant across patients, each attending to distinct sets of genes (**Supplementary Fig. 8**). To quantify the aggregate activity of these expert-associated features in each patient, we defined a Top-Gene Activity (TGA) score as a gating-weight-normalized expression score over top-ranked genes (see Methods for details). Temporal profiles of the TGA revealed divergent trajectories in cases with severe/serious and non-severe symptoms. For Expert 1, severe/serious patients exhibited progressively decreasing scores, while Expert 2 showed the opposite trend (**Fig. 4E**). These opposing trends distinguished severity-associated immune programs within this cohort. These trajectories quantify relative shifts in expert-associated activity between severity groups within a shared inflammatory disease background.

To assess cohort-level relevance, we correlated TGA with 14 clinical features across patients. The scores were most strongly associated with LDH, CRP, and Heart condition, which are biomarkers of tissue damage and systemic inflammation in COVID-19^27^, suggesting that MEGA-ODE-derived regulatory activity is associated with disease burden in this cohort (**Fig. 4F**).

To link expert-informed signals to broader biological functions and computational follow-up hypotheses, we conducted pathway and drug-signature enrichment analyses on MEGA-ODE-inferred gene-importance scores. At the gene-set level, attention-ranked genes showed enrichment for immune-regulatory programs, including JAK-STAT signaling and TNF-related responses (**Fig. 4G, Supplementary Fig. 9**); gating-score-derived gene sets showed related GO enrichments (**Supplementary Fig. 10**), consistent with established immunopathology in COVID-19^28,29^. These enrichments characterize pathway- and gene-set-level associations, while individual top-ranked genes provide hypotheses for follow-up.

Finally, we queried top-ranked genes against curated drug-target databases (details are provided in **Methods**) to identify computationally enriched compounds. Several compounds that modulate host immune or stress responses were enriched, including Nutlin-3 (an MDM2 inhibitor that activates p53), Indirubin-3’-monoxime, and the EGFR inhibitor dacomitinib (**Fig. 4H**)^30,31^. These enrichment results nominate drug signatures derived from MEGA-ODE-ranked genes for follow-up testing.

These results suggest that MEGA-ODE can model temporal expression dynamics and support retrospective patient-state analysis, while producing disease-associated pathway and drug-signature hypotheses for follow-up validation.

### MEGA-ODE prioritizes expert-associated programs during stem-cell differentiation

We next asked whether MEGA-ODE can prioritize modular expert-associated programs during cellular differentiation. We analyzed the GSE279710 stem-cell differentiation atlas, which profiles BMP4-, XAV- and Dorsomorphin-induced differentiation trajectories across multiple lineage contexts^32^. Pseudobulk profiles were constructed by aggregating cells within ordered lineage stages for each perturbation. In the BMP4 trajectory, MEGA-ODE predicted held-out expression states with high concordance (Pearson correlation = 0.951, cosine similarity = 0.910, *R*^2^ = 0.866; **Fig. 5A**), supporting the use of this setting to interrogate model-derived expert-associated programs.

**Fig. 5.**
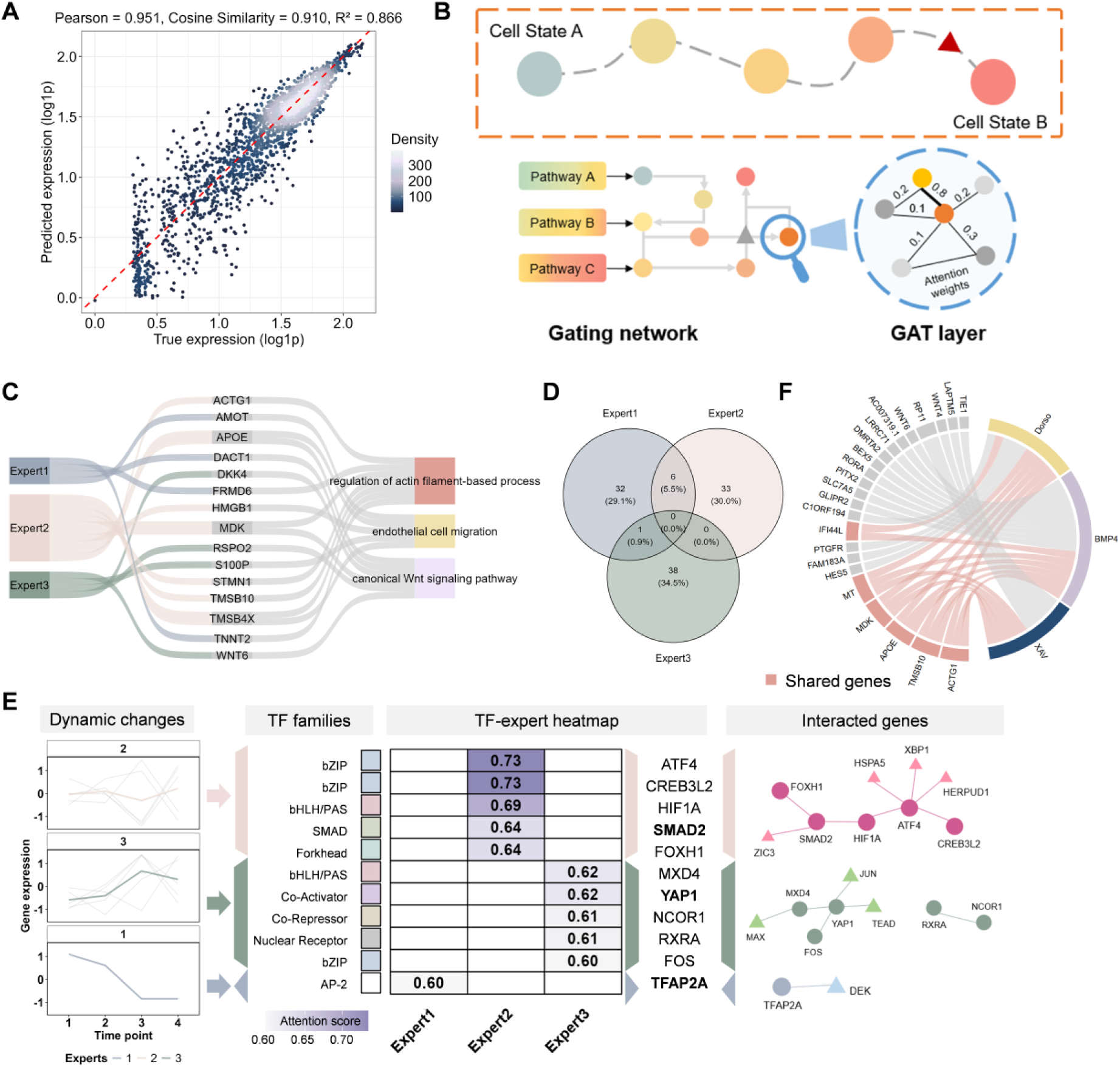
Expert-associated program prioritization during iPSC differentiation. (A) Prediction accuracy in the BMP4 differentiation trajectory. (B) Schematic of expert-level routing by the gating network and edge-level prioritization by graph attention. (C) Association between expert modules, prioritized genes, and enriched biological processes. (D) Overlap of top-ranked attention genes across experts in the BMP4 perturbation. (E) Temporal dynamics and TF-family organization of highly weighted regulators, with selected interacted genes shown for representative TFs. (F) Shared prioritized genes across BMP4, XAV and Dorsomorphin perturbation trajectories.

The mixture-of-experts architecture represented the ordered differentiation-stage profiles through partially distinct expert-associated programs. Gating weights assigned genes to expert modules, whereas graph attention weights prioritized candidate interactions within each module (**Fig. 5B, C**). In the BMP4 perturbation, attention scores were non-uniform across top- and bottom-ranked genes (**Supplementary Fig. 11**), and the overlap of highly weighted genes across experts was limited: no gene was shared by all three experts, and the largest pairwise overlap involved six genes (**Fig. 5D**). This pattern suggests that MEGA-ODE uses expert modules to represent complementary aspects of the differentiation trajectory across multiple programs.

The expert-specific programs aligned with known developmental biology and highlighted additional candidate associations. Expert 2 prioritized TF families and regulators linked to mesendoderm and stress-responsive differentiation, including bZIP, bHLH/PAS, SMAD and Forkhead factors. Among the highly weighted interactions were SMAD2-FOXH1, consistent with SMAD2/3 and FOXH1 roles in human embryonic stem-cell signaling^33^, and ATF4-CREB3L2, a less-characterized association; the SMAD2-FOXH1 interpretation is further supported by FOXH1-directed TGF-beta signaling studies^34^.

Expert 3 emphasized regulators connected to transcriptional state plasticity, enhancer-mediated transcriptional modulation and chromatin-linked signaling, including YAP1, FOS, TFAP2A, NCOR1 and RXRA (**Fig. 5E**). This organization is consistent with established YAP/TEAD transcriptional regulation and enhancer-level cooperation between YAP/TAZ/TEAD and AP-1 factors^35,36,37^; the TFAP2A-associated module highlighted a less-characterized candidate program.

We further evaluated whether selected MEGA-ODE-prioritized TF-target links were supported by independent chromatin binding evidence. Using ChIP-Atlas H9 hESC ChIP-seq datasets for FOXH1, SMAD2, YAP1 and FOS^41^, we identified nine promoter-overlap-supported relationships among the prioritized TF-target list (**Supplementary Table 1, Validated_TF_target_list sheet**). These included the known FOXH1-SMAD2 axis; four SMAD2-associated target genes (BAMBI, FOXP1, POU5F1 and TGFB1); and four YAP1-associated target genes (AJUBA, AMOTL2, TEAD1 and WTIP). These promoter overlaps provide orthogonal binding support for the highlighted TF-target links. The selected links therefore define candidates for perturbation-based tests of regulatory direction and context specificity.

Across perturbation contexts, MEGA-ODE also prioritized shared functional programs. Genes such as *IFI44L*, metallothionein genes, *MDK* and *APOE* were repeatedly highlighted across BMP4, XAV and Dorsomorphin trajectories (**Fig. 5F**). Perturbation-specific enrichment analysis linked XAV to metabolic and post-transcriptional programs, BMP4 to TGF-beta-associated developmental signaling, and Dorsomorphin to cytoskeletal organization and muscle-like differentiation programs (**Supplementary Fig. 12**). A comparison with SCENIC showed limited overlap between MEGA-ODE-prioritized TFs and regulons inferred from static co-expression and motif information (**Supplementary Fig. 13**)^38^. This difference is consistent with the two methods emphasizing different signals: SCENIC prioritizes cell-identity regulons, whereas MEGA-ODE prioritizes TFs and edges associated with time-resolved state transitions.

These analyses suggest that MEGA-ODE can predict differentiation-associated expression dynamics while prioritizing expert-specific, biologically annotated programs. The ChIP-seq overlaps provide auxiliary support for selected TF-target relationships and motivate experimental follow-up of the prioritized modules.

### MEGA-ODE enables in silico prioritization of differentiation regulators

One application of dynamical models is to computationally infer how perturbations may alter cellular trajectories. We tested whether MEGA-ODE can serve as an in silico screening tool using a chemically induced human embryonic stem cell (hESC) differentiation dataset in which cells progress toward definitive endoderm (DE) over 96 hours (GSE75748)^39^. We used the early phase (12-36 h) as observed input and used the 72- and 96-h profiles as terminal reference states for evaluating simulated trajectories. We then screened transcription factors (TFs) whose virtual overexpression was predicted to move expression profiles toward the terminal DE reference signature (**Fig. 6A**).

**Fig. 6.**
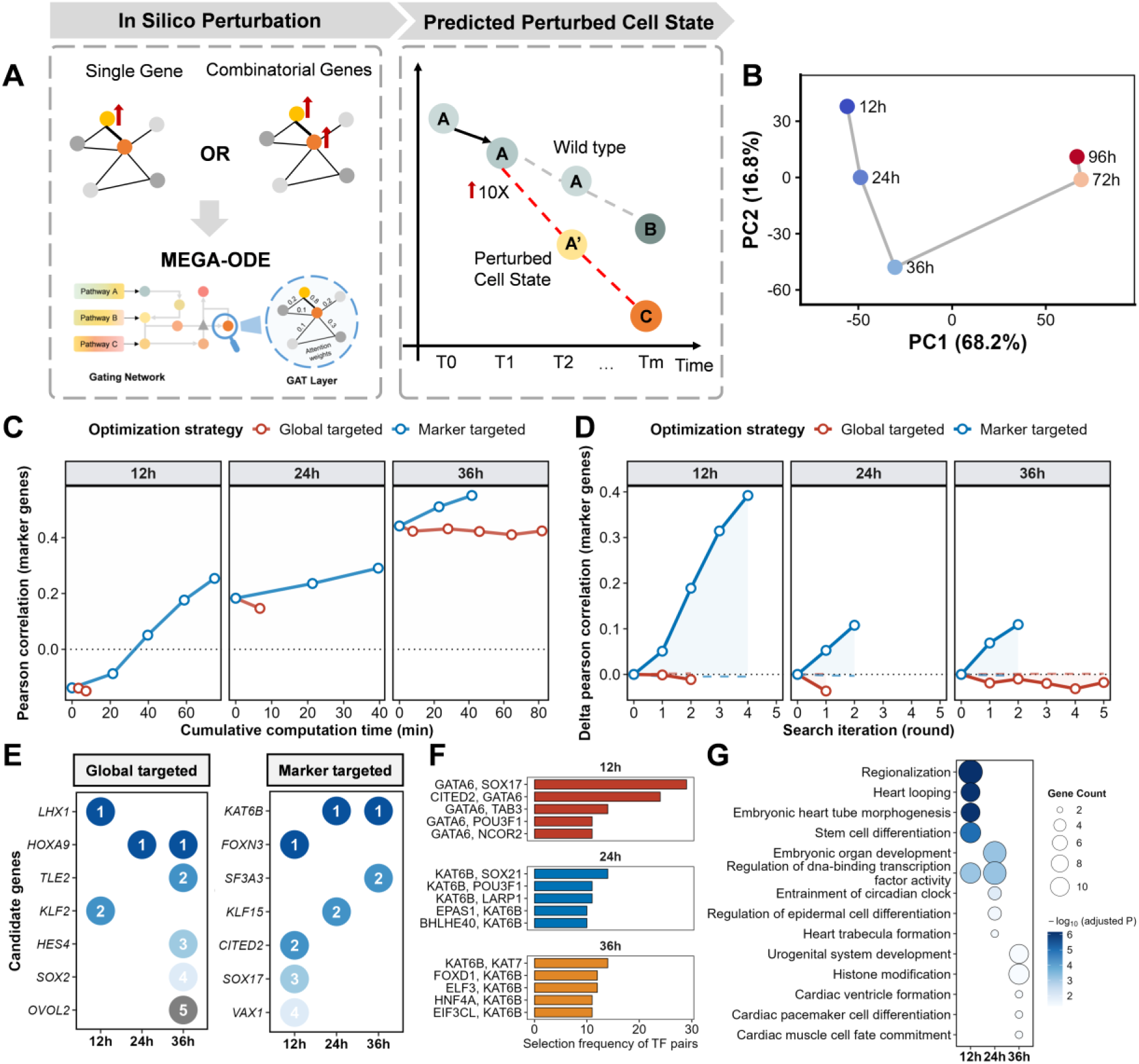
*In silico* perturbation screening for regulators of hESC-to-definitive-endoderm differentiation. (A) Schematic of the virtual perturbation framework. Single or combinatorial TF overexpression is applied at an intermediate time point (12, 24, or 36 h), and MEGA-ODE predicts the trajectory to 96 h. (B) PCA of the unperturbed time series shows progressive divergence of early (12-36 h) and late (72-96 h) states. (C) Marker Pearson score as a function of cumulative computation time for marker-targeted and global-targeted optimization. (D) Incremental improvement in marker Pearson relative to the unperturbed baseline across iterative rounds of greedy TF selection. (E) Top TFs selected by marker-targeted and global-targeted strategies at each perturbation time point. Numbers indicate the recruitment round. (F) Most frequently co-occurring TF pairs from combinatorial perturbation sets assembled by greedy search. (G) Gene Ontology biological process enrichment of combinatorial TF sets, showing temporal stratification from early lineage commitment (12 h) to later differentiation-associated annotations; heart and urogenital terms at 36 h are computational annotations requiring follow-up.

The screening space comprised 869 TF-encoding genes obtained by intersecting 5,729 highly variable genes (HVGs) with a curated set of 2,765 human TF annotations. Ten DE marker genes from MSigDB (CDH1, CXCR4, EOMES, FOXA2, GATA4, GATA6, GSC, MIXL1, SOX17, TCF7L2) served as the target signature. For each candidate TF-encoding gene, we computationally scaled its expression by 10× at an intermediate time point (12, 24 or 36 h), simulated the perturbed trajectory forward to 96 h with MEGA-ODE, and scored similarity between the predicted and reference DE transcriptome. We used marker Pearson and global Pearson in two distinct roles. During search, one of the two scores served as the objective that selected the next virtual perturbation: marker Pearson was computed on the ten DE marker genes, whereas global Pearson was computed across all 5,729 HVGs. After search, both scores were used as evaluation readouts for the trajectories produced by each objective. **Fig. 6C** reports the marker-Pearson readout for marker-targeted and global-targeted searches, directly testing how each search strategy affects the DE-marker signature. **Supplementary Fig. 14A** reports the corresponding global-Pearson readout across all HVGs, testing whether the same trajectories also remain close to the terminal transcriptome at genome scale. We then used marker Pearson as the primary ranking objective for single-TF perturbation and retained global Pearson as a complementary transcriptome-wide check.

Principal component analysis of the unperturbed time series revealed progressive transcriptional divergence, with cells at 12, 24 and 36 h forming distinct clusters that were well separated from the 72- and 96-h states (**Fig. 6B**). Pearson correlations between intermediate and terminal states were moderate (PCC(12 h, 96 h) = 0.665; PCC(24 h, 96 h) = 0.715; PCC(36 h, 96 h) = 0.779), confirming that substantial trajectory distance remained to be traversed at each perturbation window and justifying the *in silico* screen.

We applied a greedy iterative search: each round selected the TF whose virtual overexpression most increased the marker Pearson score, and the perturbed state became the new baseline for the next round. **Fig. 6C** summarizes marker Pearson as a function of cumulative computation time, whereas **Fig. 6D** shows the iterative marker-Pearson gain relative to the unperturbed baseline. The marker-targeted search produced higher marker-Pearson gains than the global-targeted search in this readout. The number of selected TFs and rounds followed the greedy stopping rule. Perturbations initiated at 12 hours yielded the largest incremental gain in PCC (perturbed minus unperturbed), even though the absolute PCC at 36 hours was higher because the starting state was already closer to DE. Within the tested window, the model assigned the largest perturbation-associated gain to the 12 h starting state. Because each greedy-search round uses a model-predicted state as the input for the next simulation, later rounds may move progressively farther from the observed training data. Predictive uncertainty and distance from the training data could accompany future iterative screens and inform experimental testing.

The top-ranked TFs emerging from this screen displayed coherent biological themes (**Fig. 6E**). Under global-targeted optimization, LHX1, a known upstream regulator associated with anterior primitive streak and DE differentiation, was selected in the first round at the 12-h window. HOXA9, a hematopoietic regulator whose overexpression from day 4 or later has been reported in hESC hematopoietic differentiation studies to promote hematopoiesis^40^, was also identified. Under marker-targeted optimization, SOX17, a core DE marker, appeared at 12 h. KAT6B, a histone acetyltransferase associated with chromatin regulation, and CITED2, a hypoxia-responsive factor, were also recovered. The latter finding is consistent with the original study’s observation that hypoxia promotes hESC-to-DE differentiation via CXCR4^39^; **Supplementary Fig. 14E** also shows that a global-targeted search nominates CXCR4, lending convergent support to the computational ranking. We classified the recovered TFs into cell-specific differentiation regulators (SOX17, CITED2 and HOXA9) and broader transcriptional modulators (KAT6B, KLF family members and TLE2). Together, these results suggest that MEGA-ODE can computationally nominate both canonical lineage-associated factors and context-specific cofactors from observational data.

Next, we performed combinatorial in silico perturbations to probe cooperative expert-associated programs. We sampled TF sets (5-10 members each) from top-ranked single-TF candidates, scored each set by the marker Pearson score and extracted the most frequently co-occurring pairs across high-scoring combinations (**Fig. 6F; Supplementary Fig. 14D**). Frequent pairs exhibited temporal stratification. At 12 h, GATA6-SOX17 co-occurred most often, consistent with early hESC-to-DE commitment. At 24 h, KAT6B paired with POU3F1, annotated as chromatin and transcriptional regulation, and with EPAS1, annotated as hypoxia signaling. At 36 h, FOXD1-KAT6B and HNF4A-KAT6B appeared among frequent co-occurrences and were annotated as later differentiation-associated nominations. These recurrent co-occurrences nominate candidate TF combinations for experimental testing. Together, this temporal layering suggests that MEGA-ODE prioritizes functionally distinct waves of transcriptional control, with early waves associated with lineage commitment and later waves shifting toward maturation.

Gene Ontology biological process enrichment of the combinatorial TF sets reinforced this interpretation (**Fig. 6G**). The 12-hour set was enriched for “regionalization”, “stem cell differentiation”, and “regulation of DNA-binding transcription”. The 24-hour set showed enrichment for “regulation of epidermal cell differentiation” and “regulation of organ development”. The 36-hour set was associated with “urogenital system development”, “cardiac ventricle formation”, and “cardiac pacemaker cell differentiation”. These 36-h annotations extend beyond the endoderm-directed protocol and may reflect broad functional annotations or mixed-lineage signals in the prioritized TF sets.

Sensitivity analyses indicated that the core rankings were stable across the tested perturbation magnitudes. At 12 h, SOX17 and KAT6B were selected across multiple 12-h perturbation magnitudes (5×, 10× and 20× virtual overexpression; Supplementary Fig. 14B, in which columns denote the nine time-by-magnitude conditions, white cells denote non-selection and colored cells encode the greedy-search selection round). **Supplementary Fig. 14C** compares marker-level and genome-level readouts, including marker Pearson, global Pearson, marker *R*^2^ and global *R*^2^, and showed higher marker-level scores for marker-targeted searches than for global-targeted searches.

Overall, these analyses show that a trained MEGA-ODE model can be used for model-based in silico perturbation ranking, providing hypotheses for experimental perturbation studies. By coupling dynamical extrapolation with iterative in silico screening, the framework offers a route to hypothesis generation in developmental and regenerative biology.

## Discussion

Dynamic omics methods now span perturbation-response models, continuous-time models and graph-informed neural architectures. Perturbation-response approaches such as scGen and neural optimal transport are powerful for predicting endpoint responses to interventions^7, 8^, whereas ODE-based frameworks are designed to model temporal trajectories from irregular or sparse observations^6,10,12^. Graph-informed models can incorporate molecular priors^13,14^, but they are often not evaluated simultaneously on missing features, unseen perturbations and unsampled time points. In this context, MEGA-ODE is positioned as a methods framework for reconstructing sparse dynamic omics landscapes and for asking how predicted states can be biologically interpreted. Through benchmarking and case studies across molecular, cellular and patient-derived datasets, MEGA-ODE improved selected held-out prediction tasks and generated experimentally testable hypotheses from gating and attention patterns.

The methodological contribution is the coupling of three inductive biases. Molecular-network priors constrain the state derivative to propagate through reported regulatory, physical or functional associations, reducing unconstrained trajectory fits that are hard to interpret^13,14^. The neural ODE backbone represents expression change as a continuous flow, which is useful when biological transitions occur between sampled time points^6,10,12^. The mixture-of-experts design adds capacity for heterogeneous programs while retaining a route to post hoc program annotation^17,18^. Graph attention then provides edge-level prioritization for association-level interpretation^19^.

Overall, these design choices make MEGA-ODE useful as a trajectory-reconstruction and perturbation-prioritization tool. In the MAPK task, the model reconstructed masked downstream expression and highlighted perturbation-associated edges, including MAPK3-MYC under KRAS(G12C) inhibition. In the COVID-19 patient analyses, predicted intermediate profiles provided additional features for retrospective disease-stage stratification, while expert-associated programs recovered immune and inflammatory signals consistent with known COVID-19 biology^25,26,27,28,29^. In the hESC-to-definitive-endoderm analysis, the in silico perturbation screen nominated TFs and TF combinations predicted to move expression profiles toward terminal marker signatures. The same screen highlighted less-characterized candidates, including KAT6B and EPAS1. These analyses illustrate how reconstructed trajectories and model-derived attributions can nominate perturbations and regulatory programs for experimental testing, consistent with recent calls for virtual-cell tools that simulate unobserved biological states^2^.

Several limitations remain. First, the model depends on a predefined biological graph, which may contain errors, omissions or context-mismatched edges. Although the model is robust to partial edge masking, future work could explore graph learning or refinement during training. Second, while the MoE structure improves interpretability, expert-level attributions remain indirect; integration with perturbation-aware saliency, counterfactual analysis or uncertainty estimation may strengthen confidence in biological interpretation. Third, attention weights and gating scores are learned to minimize prediction loss, not to estimate causal regulatory effects. Their biological interpretation is therefore correlational and requires experimental validation. Iterative virtual perturbation can also move simulated states away from the training distribution because each round uses a model-predicted state as the next input. This closed-loop use of MEGA-ODE makes uncertainty estimation, distance-to-training-data diagnostics and experimental validation particularly important before prioritizing high-confidence interventions. Fourth, some experts were rarely activated, raising the possibility of underused model capacity. Mechanisms for expert regularization or diversity promotion could improve modular efficiency. Finally, although we evaluate MEGA-ODE on bulk and pseudobulk data, extending it to fully single-cell resolution remains an open direction and may require tailored adaptations to address stochasticity and dropout noise.

Several extensions may strengthen MEGA-ODE’s utility. Moving beyond static graphs to dynamic or context-specific regulatory networks that evolve during differentiation or disease progression would relax dependence on fixed prior structure. Integrating chromatin accessibility, proteomic or metabolomic layers alongside transcriptomic readouts would more faithfully capture the multi-layered logic of gene regulation. In patient-derived datasets, the COVID-19 cohort provided an initial example of predicted intermediate profiles augmenting retrospective stage stratification and expert programs recovering severity-linked immune signals. Combining molecular predictions with imaging and longitudinal phenotypic data may further improve disease stratification and treatment-response forecasting, but substantially larger cohorts and prospective validation would be required before clinical use. Critical assessment of which inductive biases, including graph structure, smoothness of ODE dynamics and mixture assumptions, are most likely to fail in new contexts will be essential for guiding future applications.

Overall, MEGA-ODE provides a biologically structured continuous-time strategy for modeling sparse dynamic omics data. By linking graph-constrained ODE dynamics with expert routing and attention-based prioritization, the framework supports held-out prediction, model-derived biological interpretation and virtual-perturbation ranking within a common workflow. This integration provides a practical tool for prioritizing experimentally testable regulatory hypotheses from dynamic omics data.

## Methods

### MEGA-ODE method overview

MEGA-ODE takes two inputs: a molecular interaction graph and a time-ordered omics matrix in which rows represent samples or pseudo-bulk states and columns represent genes or proteins. The method is organized around three linked designs: graph-constrained propagation to keep the learned dynamics biologically plausible, a neural ODE backbone to represent continuous transitions between sampled states, and a mixture-of-experts gate to increase expressiveness for heterogeneous biological programs. The same architecture also returns expert weights and graph-attention scores that can be analyzed after training. We first describe the model architecture and training procedure, followed by benchmarking models, sensitivity, scalability and ablation analyses, dataset preprocessing, model interpretation and evaluation metrics.

### MEGA-ODE model architecture

MEGA-ODE integrates biological association priors, dynamic ODE modeling and expert-driven trajectory modules into a unified computational framework for reconstructing omics time series, predicting responses to unseen perturbations and prioritizing latent expert-associated programs. Inspired by dynamical systems theory, MEGA-ODE treats gene or protein expression evolution as a continuous-time flow conditioned on supplied biological interaction graphs. The overall framework includes two key components: a gating network that adaptively weights latent expert modules representing alternative dynamical patterns, and a graph-based neural ODE module that simulates continuous-time omics evolution under graph-constrained dynamics.

### Gating Network: learning context-specific expert weights

Different cellular contexts often activate distinct regulatory subcircuits, such as differentiation versus stress response programs. To capture such heterogeneity, MEGA-ODE employs a mixture-of-experts (MoE) gating mechanism, where each expert represents a distinct learned dynamical component that can later be interrogated for biological enrichment. This gating network is implemented as a three-layer multilayer perceptron (MLP) consisting of linear layers with ReLU activations. For an input vector x, the MLP outputs K gating scores, one for each expert module. These scores are normalized by a softmax function and used to weight the outputs of the corresponding experts:

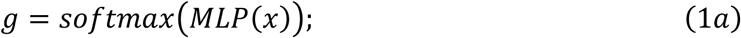

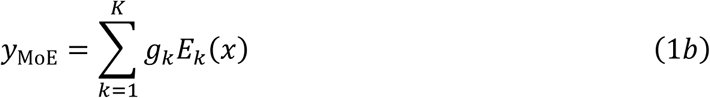

where x is the input molecular profile, *g_k_* is the normalized weight assigned to expert k, *E_k_*(*x*) is the output of expert k and K is the number of experts.

Thus, the gating module functions as a soft weighting layer over learned dynamical components; biological interpretation of these components is performed post hoc through enrichment and attention analyses.

### Graph Encoder: embedding prior biological interactions into graph-aware latent coordinates

MEGA-ODE incorporates prior biological networks (PPI, GRNs, pathways) using a Graph Attention Network (GAT)^19^. The GAT module encodes prior biological network information to obtain gene embeddings. Each gene node aggregates information from its neighbors through self-attention, enabling the model to learn the relative importance of each neighbor:

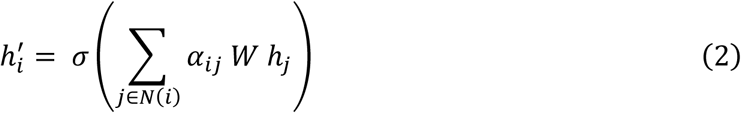

where ℎ*_j_* and ℎ*_i_*′ are the input and updated node features, respectively; *α_ij_* is the model-assigned attention weight on the edge from node j to node i; *N*(*i*) is the neighbor set of node i; W is a learnable weight matrix; and *σ*(·) is a nonlinear activation function.

The unnormalized attention score and normalized attention coefficient are computed as:

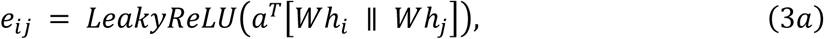

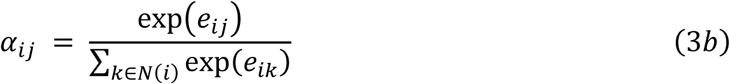

where a is the attention vector and || denotes concatenation.

The attention mechanism learns context-dependent edge weights, which we use to prioritize candidate graph edges, including TF-target-like edges when the supplied graph and dataset support that interpretation, or pathway hubs for downstream interpretation.

We employ multi-head attention to stabilize training and enhance representation power. The GAT consists of two layers, producing final embeddings for each node i. For each node i, the multi-head update is:

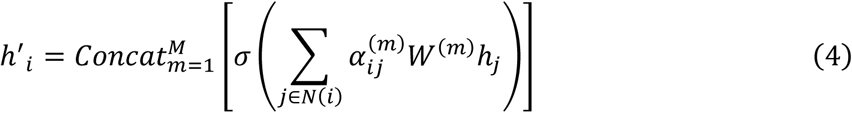

where M is the number of attention heads, concat denotes concatenation, and *W*^(*m*)^ and 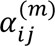 are the weight matrix and attention coefficient for head m, respectively.

Multi-head attention provides multiple learned channels for aggregating information across the prior graph. Thus, the GAT transforms raw omics measurements into a graph-aware latent space z conditioned on the supplied prior network.

### Graph-based Neural ODE: modeling continuous-time molecular dynamics

The latent representations from the GAT encoder are passed through a Neural Ordinary Differential Equation (Neural ODE) model parameterized using a GAT:

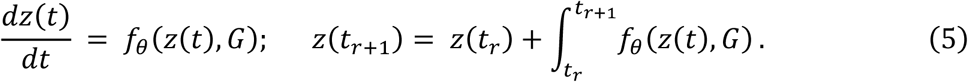

where *z*(*t*) is the graph-aware latent state at time t, G is the supplied biological graph, *f_θ_* is the GAT-parameterized ODE function and theta denotes learnable parameters. The integral was numerically solved with a fourth-order Runge-Kutta method.

The ODE models smooth temporal transitions in the latent molecular state. It captures nonlinear dynamical responses such as feedback-like patterns, differentiation trajectories or stimulus-induced expression changes. The GAT parameterization restricts information propagation in 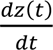 to the supplied gene-gene, protein-protein or pathway-level graph.

### Linear decoder: reconstructing observable molecular phenotypes

The ODE-evolved latent state z(t) is decoded back into gene or protein expression space using a two-layer linear decoder to predict the target profiles. The decoder acts as an observer function in dynamical systems terminology, mapping latent state trajectories to observable transcriptomic or proteomic outputs.

### Model training

#### Dataset splitting

For the **feature mask task**, the dataset was randomly divided into training, validation, and test sets with a ratio of 7:2:1. For the **drug mask task**, samples were partitioned with an 8:1:1 training, validation, and test ratio. In the **time-series prediction task**, given N temporal sampling points *t*_1_, *t*_2_, …, *t_N_*, the model was trained using data from *t*_1_ to *t_N_*_−1_ and evaluated on the final unseen time point *t_N_*.

#### Training procedures

Model training was conducted with an early stopping strategy to prevent overfitting. Hyperparameter optimization was performed using Bayesian optimization implemented with the Optuna framework, which adaptively explores the search space to identify the optimal configuration for each task. The loss function combined mean squared error (MSE) and Pearson correlation-based components to jointly capture absolute expression deviations and relative expression concordance between predictions and ground truth. Task-specific hyperparameter settings and optimization ranges are summarized in the Supplementary Table 1 (MEGA-ODE Parameters and Baselines Parameters sheets). In the feature masking task, our model generates predictions for all features. During training, only the unmasked (training) features are used to compute the loss, while during evaluation, the prediction errors on the masked (test) features are assessed.

#### Parameter settings

The mixture-of-experts layer was implemented with a hidden dimension of 128 and comprised four expert networks. Both the graph attention network encoder and the neural ordinary differential equation block (three layers of GAT) employed a hidden size of 256 with four attention heads. The decoder consisted of two fully connected linear layers with a hidden dimension of 128.

For feature mask, drug mask, and time series prediction benchmarks, models were trained with task-specific learning rates and weight decay coefficients; the exact values and optimization ranges are listed in the Supplementary Table 1 (MEGA-ODE Parameters and Baselines Parameters sheets). Across all benchmarks, early stopping based on validation performance was applied with a patience of 50 epochs. Models were trained for up to 5,000 epochs, except for drug mask benchmarking, which used up to 2,000 epochs.

### Benchmarking models

#### LSTM

We used a single-layer Long Short-Term Memory (LSTM) network as a baseline for time-series modeling. The model takes the time-series gene expression matrix as input. The hidden layer has 16 recurrent units, followed by a linear projection to predict expression at the next time point. During training, the model learns from gene expression profiles from *t*_1_ to *t_T_*_−1_ and predicts the state at *t_T_*. Multi-step forecasts are obtained by feeding predictions from the model back into itself. This discrete-time design captures short-term dependencies but cannot model continuous or irregular sampling, which MEGA-ODE addresses.

#### GLSTM

To integrate both temporal dynamics and graph topology, we implemented a Graph-LSTM (GLSTM) model. Each node represents a gene or protein, and edges denote known biological interactions between them. The model first encodes the graph structure using two GAT layers. The outputs from these layers are concatenated to form node-level embeddings. These embeddings are then processed by a single-layer LSTM with 128 hidden units to learn temporal dependencies. The final fully connected layer projects the LSTM output to predict expression values at multiple future time points.

#### MLP-based neural ODE

For the baseline Neural ODE, the ODE function f was parameterized as a four-layer multilayer perceptron (MLP) with hidden dimensions of 256-128-128 and Tanh activation functions.

#### moeLSTM, moeGLSTM, and moeNODE

Building upon the LSTM, GLSTM, and Neural ODE architectures described above, we introduced mixture-of-experts extensions (moeLSTM, moeGLSTM, and moeNODE). Each model integrates multiple expert modules with a shared gating network that adaptively weights their outputs. The gating network architectures of moeLSTM, moeGLSTM, and moeNODE follow the same design as in MEGA-ODE.

#### Random forests

A Random Forest (RF) classifier was implemented in R using the caret package. Model robustness was ensured through 5-fold cross-validation repeated three times. For patient-stage classification, all observed and augmented profiles from each patient were assigned to the same fold to avoid leakage. The number of features considered at each split (mtry) was set to the square root of the total features, a common practice. Models were evaluated across a range of tree counts (ntree = 100 and 200), and the final classifier was selected based on the highest cross-validated accuracy.

#### Sensitivity analyses

We assessed sensitivity in three places: prior-graph incompleteness by masking increasing proportions of edges, measurement noise by adding Gaussian perturbations to input features, and virtual-perturbation magnitude by repeating the hESC TF screen at 5×, 10× and 20×. These analyses tested whether predictions and regulator rankings were stable to plausible variation in graph knowledge, assay noise and computational perturbation strength, and they correspond to the robustness analyses reported in Fig. 2D, Supplementary Fig. 3 and Supplementary Fig. 14B, C.

#### Scalability analysis

Scalability was evaluated across benchmark sizes and screening spaces using observed computation time and search progression. The benchmarking tasks covered matrices ranging from hundreds of proteins to thousands of genes, and the in silico screen recorded cumulative computation time while ranking 869 TF-encoding genes across three starting time points. For the combinatorial stage, we restricted the search pool to the top 50 single-TF candidates at each time point before sampling 5-10-member sets, which kept the screen interpretable and computationally bounded while still allowing recurrent co-occurring TF pairs to be identified. These analyses correspond to the in silico screening runtime and search-progression results reported in Fig. 6C, D and F.

#### Ablation analyses

To isolate contributions of temporal modeling, graph priors and expert decomposition, we compared MEGA-ODE with LSTM, GLSTM, Neural ODE and mixture-of-experts extensions of these baselines. The recurrent and ODE baselines tested time-series modeling without the full MEGA-ODE design, graph-augmented baselines tested the effect of supplied molecular networks, and MoE variants tested whether expert decomposition improved held-out prediction and produced annotatable programs. Edge-masking experiments further ablated the completeness of the prior graph, linking the robustness analysis to the role of graph-constrained dynamics. These analyses correspond to the baseline and module-comparison results reported in Fig. 2B, C, Fig. 3C, Fig. 4B and Supplementary Figs. 2 and 3.

#### Preprocessing for L1000 dataset

The L1000 transcriptomics data used for the benchmarking experiments were collected at 1, 3, 6 and 24 h after treatment with peptides and biological agents (34 perturbation conditions), including the expression of 978 landmark genes. The gene expression matrix was log-normalized. The training, validation and test split for gene-mask prediction was 7:2:1.

#### Preprocessing for CPPA dataset

The CPPA benchmarking data contained expression profiles of 549 proteins across 422 samples. Proteins with expression values equal to zero in more than 30% of samples and samples with zero values across multiple features were excluded, leaving 374 proteins and 410 samples for modeling. The benchmark included two tasks. For the protein-mask task, we used data from the MCF7 cell line, which includes protein expression profiles collected at 4, 24 and 48 h after perturbation with nine drugs. The training, validation and test split for protein-mask prediction was 7:2:1. For the drug-mask task, after excluding samples with missing time points, samples treated with nine drugs were used for benchmarking. The training, validation and test split for drug-mask prediction was 8:1:1.

#### Dataset for KEGG pathway case study

For the MAPK pathway case study, we used the GSE103021 dataset, which contains transcriptomic profiles of NCI-H358 cells treated with ARS-1620 or Trametinib for 4, 24 and 48 h^20^. The prior network of the MAPK signaling pathway was derived from the KEGG database. The split point was defined at MAPK1 (ERK2) and MAPK3 (ERK1). ERK and upstream nodes on the KRAS-RAF-MEK-ERK axis were used as observed training nodes, whereas post-ERK downstream nodes were treated as masked targets while remaining connected in the pathway graph. The model was trained on the 4 h expression profile and evaluated on downstream target prediction at 24 and 48 h. Attention ranks were then compared between the ARS-1620 and Trametinib models separately for upstream training-node edges and downstream testing-node edges.

#### Dataset for COVID-19 time series benchmarking and case study

For benchmarking, we used the GSE151513 dataset, which contains transcriptomic profiles of human lung epithelial cells following SARS-CoV-2 infection.

For clinical stage prediction, the GSE157859 dataset was used, comprising PBMC samples obtained from 13 COVID-19 patients diagnosed with mild, moderate, or severe symptoms. Each patient was sampled at two time points: the Treatment Stage and the Convalescence Stage. In the original setting, the model directly predicted the clinical stage using gene expression data from the Treatment Stage. In the data augmentation setting, both Treatment and Convalescence Stage samples were used to train MEGA-ODE, which subsequently predicted gene expression profiles at future time points. For downstream disease-state classification, cross-validation splits were defined at the patient level: all observed and MEGA-ODE-predicted profiles from the same patient were assigned to the same fold, preventing augmented profiles from a held-out patient from entering classifier training.

For the GSE212041 dataset, we filtered samples to include only those collected on days 0, 3 and 7 after infection, resulting in a total of 85 samples for analysis. All datasets were log-normalized, and the top 3,000 highly variable genes were selected for downstream analysis.

#### Dataset for cell differentiation analysis

For cell differentiation analysis, we used the GSE279710 dataset, which contains single-cell transcriptomic profiles obtained under BMP4, XAV and Dorsomorphin perturbations^32^. To construct pseudo-bulk expression profiles, the mean expression of each gene was calculated across cells belonging to the same cell type. All datasets were log-normalized, and the top 3,000 highly variable genes were selected for downstream analysis.

For dynamic modeling, pseudo-bulk profiles were organized as ordered lineage-stage inputs by aggregating specific cell types selected to approximate consecutive differentiation stages. For the BMP4-perturbed dataset, we selected Mesendoderm, Definitive endoderm, Foregut progenitor, and Pancreas cells. For the XAV-perturbed dataset, we selected Epiblast, Mesendoderm, First heart field ventricular cardiomyocyte, and Second heart field ventricular cardiomyocyte cells. For the Dorsomorphin-perturbed dataset, we selected Epiblast, Mesendoderm, Ventral neural tube, and Anterior neural plate cells.

#### ChIP-seq support for prioritized TF-target relationships

To provide orthogonal binding evidence for selected TF-target relationships prioritized in the pluripotent-stem-cell differentiation analysis, we queried ChIP-Atlas peak sets generated from H9 hESC samples^41^. We focused on four transcriptional regulators highlighted in the MEGA-ODE TF-target analysis: FOS, FOXH1, SMAD2 and YAP1. ChIP-seq peaks were downloaded using a stringent threshold of Q < 1 × 10^-5^. For each prioritized TF-target pair, target-gene promoter intervals were mapped to the genome build used by the ChIP-seq peak set, and an interaction was considered ChIP-supported if at least one TF ChIP-seq peak overlapped a promoter interval of the corresponding target gene. This analysis identified nine promoter-overlap-supported relationships among the prioritized list: FOXH1-SMAD2; SMAD2-BAMBI, SMAD2-FOXP1, SMAD2-POU5F1 and SMAD2-TGFB1; and YAP1-AJUBA, YAP1-AMOTL2, YAP1-TEAD1 and YAP1-WTIP.

#### Dataset for in silico perturbation screening

For the hESC differentiation analysis, we used the GSE75748 dataset, which profiled human embryonic stem cells differentiating toward definitive endoderm by single-cell RNA-seq at 0, 12, 24, 36, 72 and 96 h^39^. We used the processed expression matrix as time-ordered profiles, log-normalized the data and retained 5,729 highly variable genes for the perturbation screen. For model training, the 12-36 h time window was used as the training set, and extrapolation performance was evaluated on the 72 and 96 h time points. For in silico perturbation, the trained model was used to predict trajectories after computationally scaling the expression of target genes at 12, 24 or 36 h. The 5×, 10× and 20× settings were treated as sensitivity analyses, not experimentally optimized doses.

### Model interpretation and biological analysis

#### Biological prior information

We used two sources of prior information for graph construction: protein-protein interactions (PPI) from the STRING database (https://string-db.org/)^42^ and gene-gene interactions (functional gene network) from HumanNet v3 (https://www.inetbio.org/humannet/)^43^. For a protein-based time-series expression matrix, we used the PPI as prior information. For a gene-expression matrix, we used the gene-gene network from HumanNet to represent prior information.

#### Bioinformatics analysis

We used clusterProfiler to perform Gene Ontology (GO) enrichment and Gene Set Enrichment Analysis (GSEA)^44^. We also used Enrichr with LINCS_L1000_Chem_Pert_up and LINCS_L1000_Chem_Pert_down signatures to perform drug set enrichment, which identified small molecules computationally associated with the target-gene signatures. For the pluripotent-stem-cell differentiation case study, SCENIC was used as a regulon-based comparator for transcription factor prioritization^38^.

#### Top-Gene Activity

We used Top-Gene Activity to quantify the activity level of a selected feature set in each patient as a gating-weight-normalized expression score.

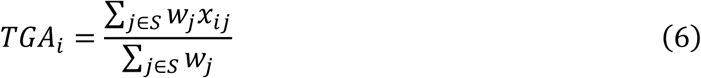

where S is the selected gene set, *x_ij_*is the expression of gene j in patient i and *w_j_* is the maximum gating weight of gene j across patients. The denominator normalizes the score by the total weight assigned to the selected genes.

#### In silico perturbation screening

To identify candidate regulators predicted to move expression profiles toward a differentiation target signature, we implemented an in silico perturbation pipeline on top of the trained MEGA-ODE model. The hESC-to-definitive-endoderm differentiation dataset (GSE75748)^39^ comprises single-cell RNA-seq measurements collected at 0, 12, 24, 36, 72 and 96 h. MEGA-ODE was trained on the early time window (12-36 h) and evaluated on its ability to extrapolate to 72 and 96 h.

#### Screening space and target signature

The screening space comprised 869 transcription factors (TFs) obtained by intersecting the 5,729 highly variable genes with a curated set of 2,765 human TFs^45^. The target definitive endoderm signature consisted of ten marker genes from MSigDB: *CDH1*, *CXCR4*, *EOMES*, *FOXA2*, *GATA4*, *GATA6*, *GSC*, *MIXL1*, *SOX17*, and *TCF7L2*.

#### Single-TF perturbation with iterative greedy search

For each candidate TF, we virtually overexpressed it by scaling its expression by 10× in the primary screen at an intermediate time point (12, 24, or 36 h). The modified expression profile was fed into MEGA-ODE, which predicted the resulting trajectory from the perturbation time point forward to 96 h. The outcome was scored using two objectives: a marker-targeted objective that computed the Pearson correlation between the predicted and observed 96-hour expression profiles across the ten DE markers, and a global-targeted objective that computed the Pearson correlation across all 5,729 HVGs. The marker-targeted objective was used as the primary ranking criterion because single-gene perturbations produce localized effects that are better captured by a focused signature than by a genome-wide metric, whereas whole-transcriptome similarity tends to dilute marker-level gains.

To identify sets of TFs that jointly improve differentiation, we implemented a greedy iterative search. In each round, the TF whose overexpression most increased the marker Pearson score was selected, and the perturbed state became the new baseline for the next round. This procedure was repeated until no further improvement was observed. The iteration number at which each TF was selected was recorded to characterize the order of recruitment.

For sensitivity analysis, we additionally repeated the single-TF overexpression screen at 5× and 20× perturbation magnitudes. Together with the primary 10× screen, these analyses tested whether the top-priority regulators remained stable across 5×, 10× and 20× virtual overexpression strengths. These auxiliary screens were evaluated using the same marker-targeted and global-targeted objectives as the primary analysis.

#### Combinatorial TF perturbation

To probe cooperative expert-associated programs, we performed combinatorial *in silico* perturbations in two stages. First, the single-TF screen provided a ranked list of candidate regulators for each starting time point. Second, we sampled TF combinations of 5-10 members from the top-ranked candidates, using the top 50 TFs from the first-round screen as the search pool for each time point, and evaluated each set by the marker-targeted Pearson score at 96 h. Sets were constructed independently for each starting time point (12, 24, and 36 h). From the ensemble of high-scoring sets, we extracted the most frequently co-occurring TF pairs and ranked them by their occurrence frequency.

To determine whether additional search rounds were warranted, we monitored a composite improvement score that integrated Pearson correlation, cosine similarity, coefficient of determination (R^2^), MSE and Euclidean reconstruction error between the predicted and observed terminal states. Search was terminated when newly proposed perturbation sets produced only marginal improvement in this aggregate score relative to the current best state, indicating that the search had reached a practical plateau. Because the biological interpretation in the main text is based on the marker-directed objective, the marker Pearson score was retained as the ranking statistic for reporting top combinations and co-occurring TF pairs.

#### Gene Ontology enrichment

Gene Ontology (GO) biological process enrichment was performed on the combinatorial TF sets using clusterProfiler. Enrichment significance was assessed using Benjamini-Hochberg-adjusted P values.

#### Evaluation Metrics

We evaluated regression and classification performance using Pearson correlation, cosine similarity, coefficient of determination (R^2^), mean squared error (MSE), mean absolute error (MAE), accuracy (ACC) and F1-score.

#### Regression metrics

For molecular-state prediction, reconstruction and trajectory-forecasting tasks, regression performance was evaluated using Pearson correlation, MSE and MAE as the principal metrics, with cosine similarity and the coefficient of determination R² reported as complementary measures.

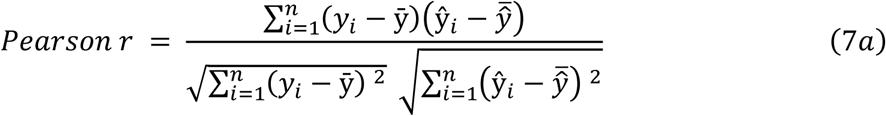

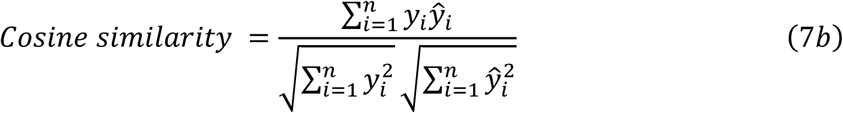

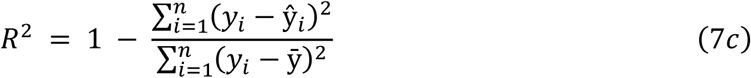

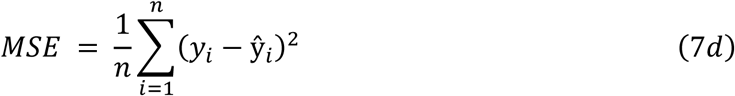

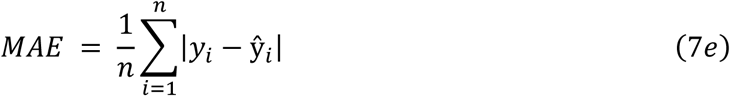

Here, *y_i_* and *̂Y_i_* denote the observed and predicted values for sample i, *̄y* and 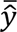 denote their means, and n is the number of evaluated samples. Pearson correlation and cosine similarity quantify linear and angular agreement; R^2^measures the fraction of variance explained; MSE and MAE quantify squared and absolute prediction errors.

#### Classification metrics

For the COVID-19 disease classification task, classification performance was evaluated primarily using accuracy and F1-score, with precision and recall reported as complementary metrics.

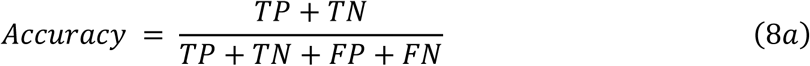

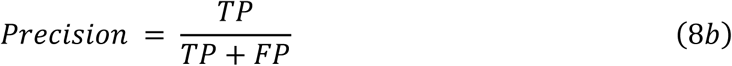

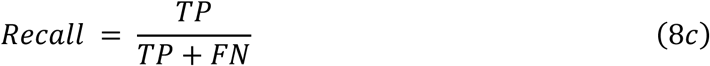

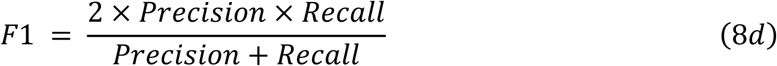

where TP, TN, FP and FN refer to true positives, true negatives, false positives and false negatives, respectively. Accuracy measures the overall proportion of correctly classified samples, whereas precision, recall and F1 describe positive predictive value, sensitivity and their harmonic mean under class imbalance.

## Data availability

All datasets analyzed in this study are publicly available from the resources cited in the manuscript, including L1000, the Cancer Perturbed Proteomics Atlas (CPPA), GEO accessions GSE103021, GSE151513, GSE157859, GSE212041, GSE279710 and GSE75748, and ChIP-Atlas. Dataset inventory, preprocessing context and associated task usage are summarized in the Supplementary Table workbook (Datasets sheet).

## Code availability

The code for MEGA-ODE is available at https://github.com/Candlelight-XYJ/MEGA-ODE and a demo is available at http://bio2info.top/MEGA-ODE/.

## Supplementary information

Supplementary information, including Supplementary Figs. 1–14 and Supplementary Table 1, is available with this paper.

## Author Contributions

P.Z., L.X., F.H., H.W. and Y.L. conceived and supervised the study. Y.X. and Y.L. developed and implemented the MEGA-ODE model. Y.X. conducted the computational experiments and benchmark evaluations, performed the downstream biological analyses and prepared the figures. Y.X., Y.L. and C.T. analysed and interpreted the results. R.G. contributed to figure development. All authors contributed to the writing and revision of the manuscript and approved the final version.

## Acknowledgements

The work was supported by the National Key Research and Development Program of China (2025YFA1309400) and the Science and Technology Commission of Shanghai Municipality (STCSM; Grant No. 25JS2850100). P.Z. acknowledges support from the NSFC (grant nos. 12288101, 8206100646, and T2321001), the Fundamental and Interdisciplinary Disciplines Breakthrough Plan of the Ministry of Education of China (grant no. JYB2025XDXM502), and the Fundamental Research Funds for the Central Universities.

