## Supplementary Materials for "MEGA-ODE: Learning Biologically Structured and Navigable Continuous Perturbation Dynamics from Sparse Omics"

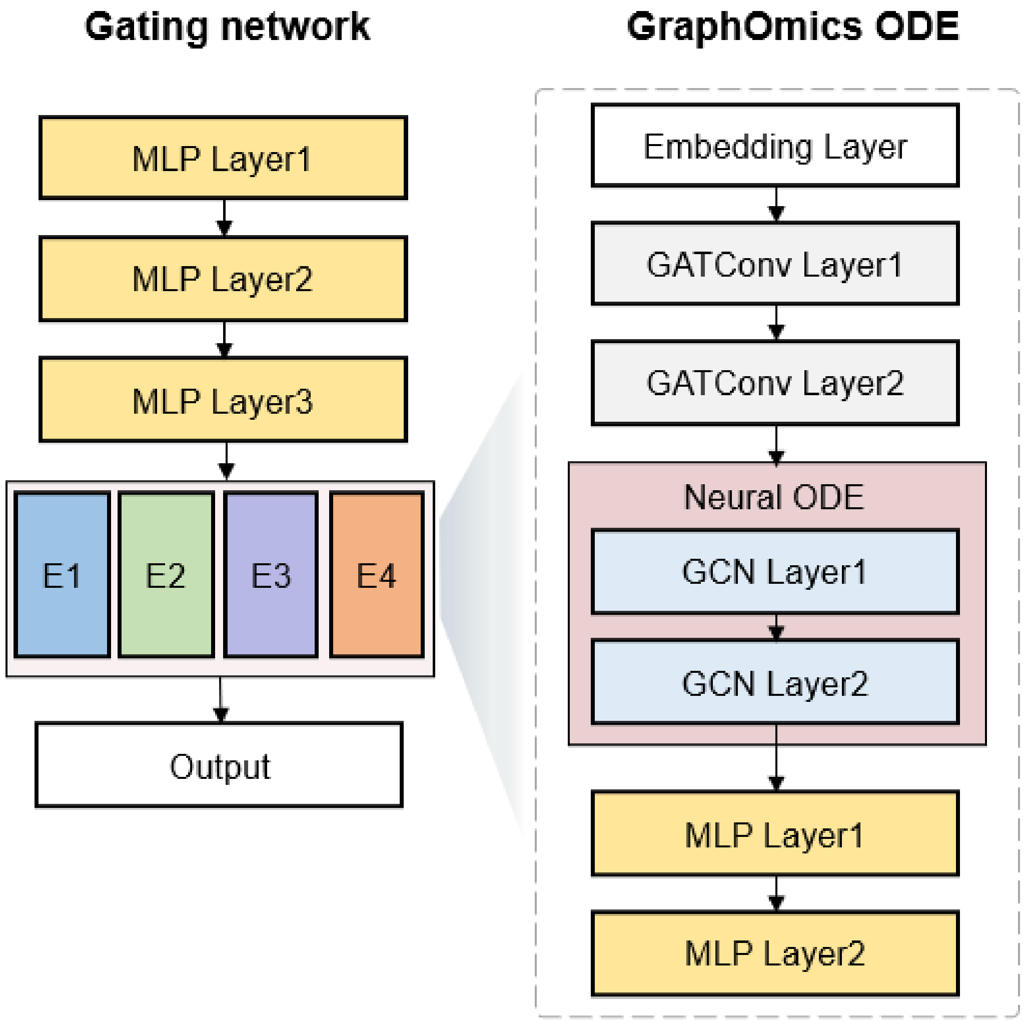


Supplementary Fig. 1 Model architecture of MEGA-ODE. MEGA-ODE consists of a gating network and a graph-based neural ODE block.

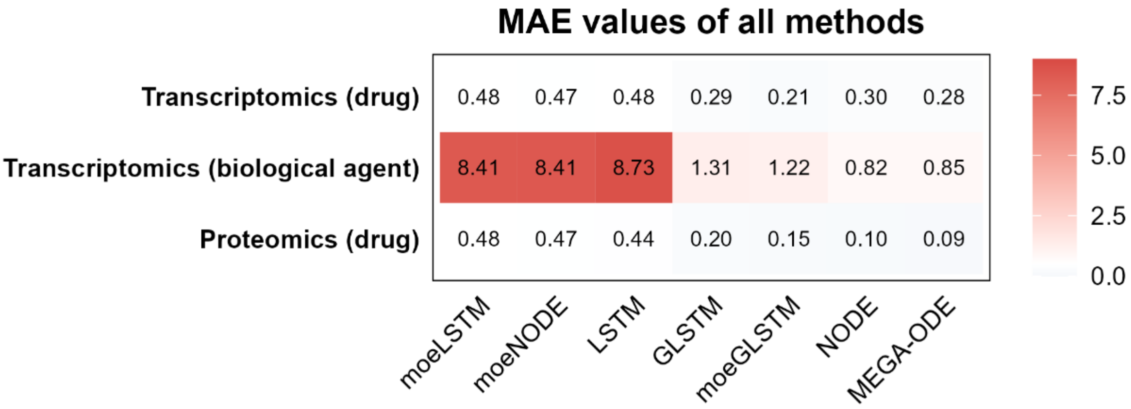


Supplementary Fig. 2 MAE results for feature-mask benchmarking.

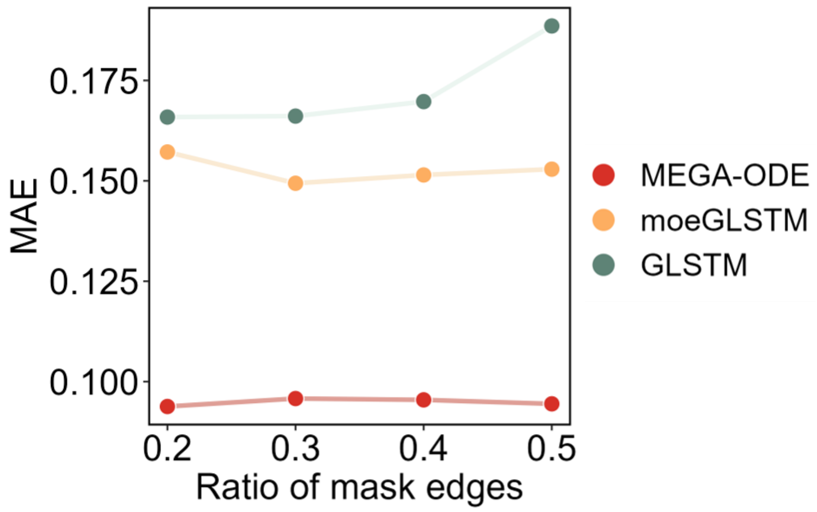


Supplementary Fig. 3 Performance of models under different edge-masking ratios.

| Drug | MOA | MOA Description |
| --- | --- | --- |
| AZD8055 | Metabolic regulation | Serine/threonine-protein kinase mTOR |
| DACTOLISIB | Metabolic regulation | Serine/threonine-protein kinase mTOR; Phosphatidylinositol 4,5-bisphosphate 3-kinase catalytic subunit gamma isoform |
| GSK1838705A | Tyrosine kinase signaling | \ |
| DASATINIB | Tyrosine kinase signaling | Tyrosine-protein kinase ABL1, Lck, Yes, Fyn |
| GEFITINIB | Tyrosine kinase signaling | Epidermal growth factor receptor |
| GSK690693 | PI3K-AKT signaling | RAC-gamma serine/threonine-protein kinase |
| LAPATINIB | Tyrosine kinase signaling | Epidermal growth factor receptor; Receptor tyrosine-protein kinase erbB-2; Eukaryotic elongation factor 2 kinase |
| PD173074 | Tyrosine kinase signaling | \ |
| TRAMETINIB | Cell cycle regulation | Dual specificity mitogen-activated protein kinase kinase 1; Dual specificity mitogen-activated protein kinase kinase 2 |

Supplementary Fig. 4 Drugs used for training and testing in the drug-mask task.

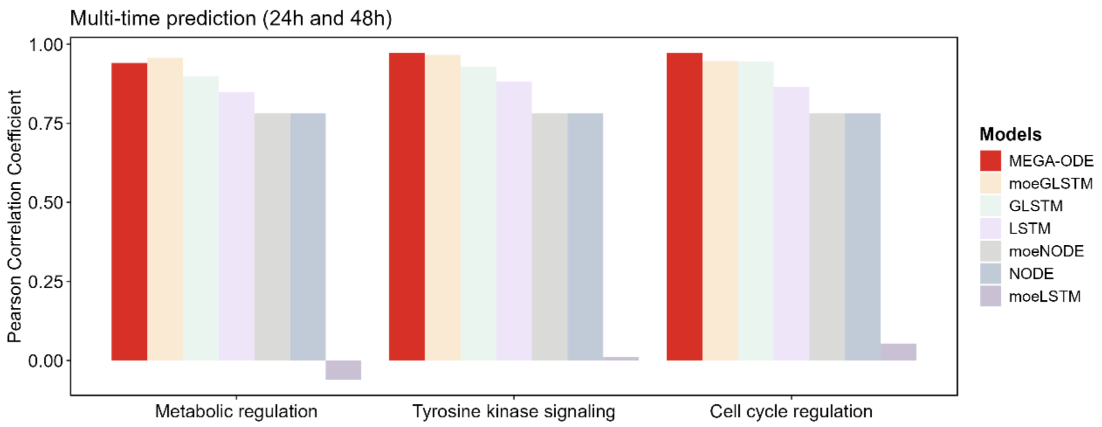


Supplementary Fig. 5 Cross-MOA generalization performance. Models were trained and validated on two mechanisms of action (MOAs) and evaluated on an unseen third MOA. The x-axis denotes the held-out MOA, and the y-axis shows the Pearson correlation coefficient between predicted and observed protein expression.

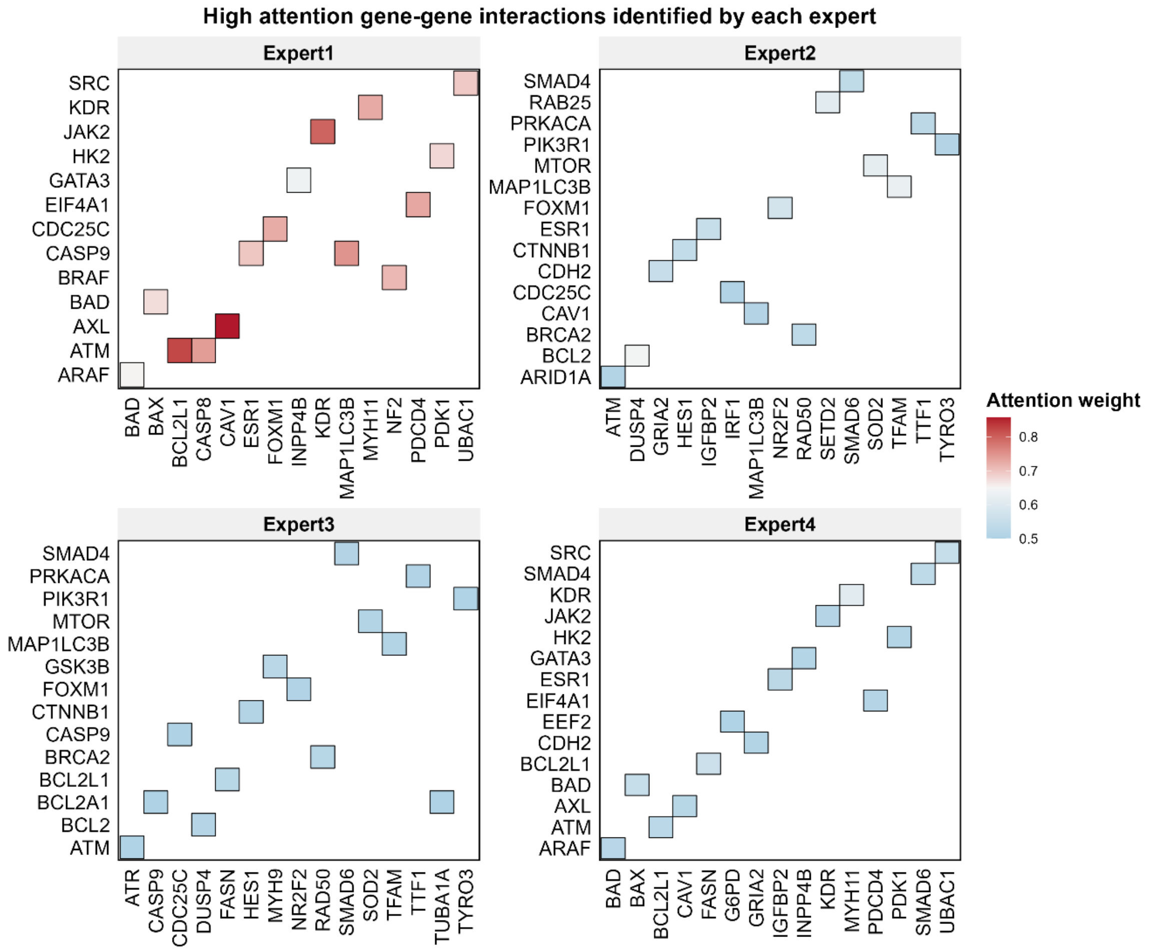


Supplementary Fig. 6 Heatmap of top-ranked attention edges across four expert modules.

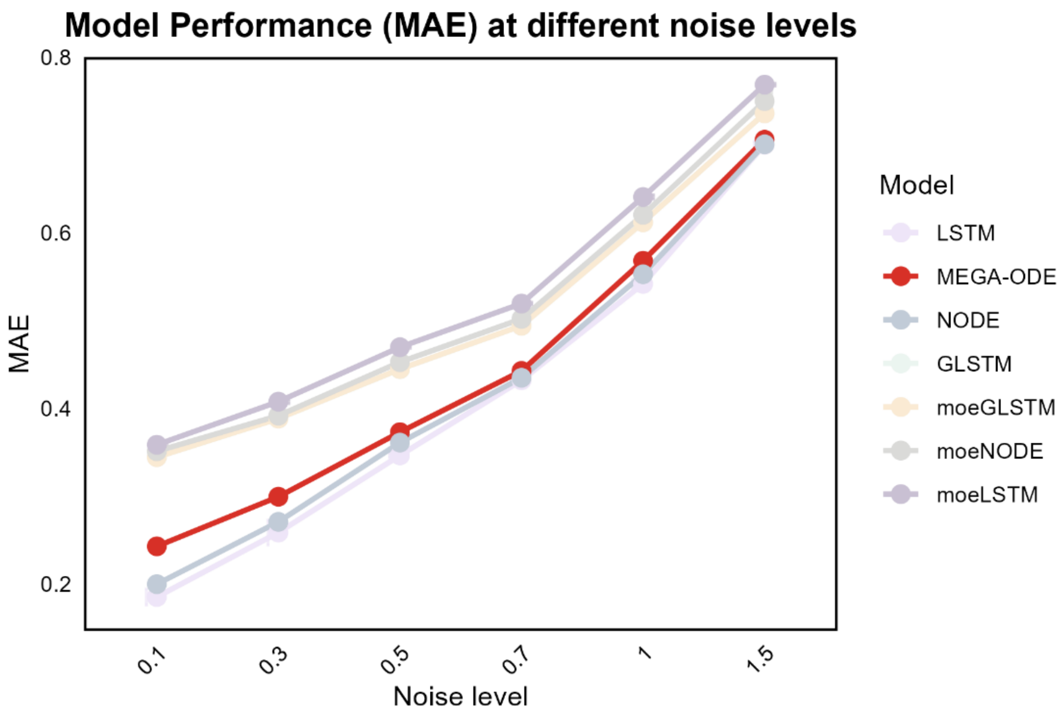


Supplementary Fig. 7 Model performance (MAE) at different noise levels for the unseen-time-point prediction task.

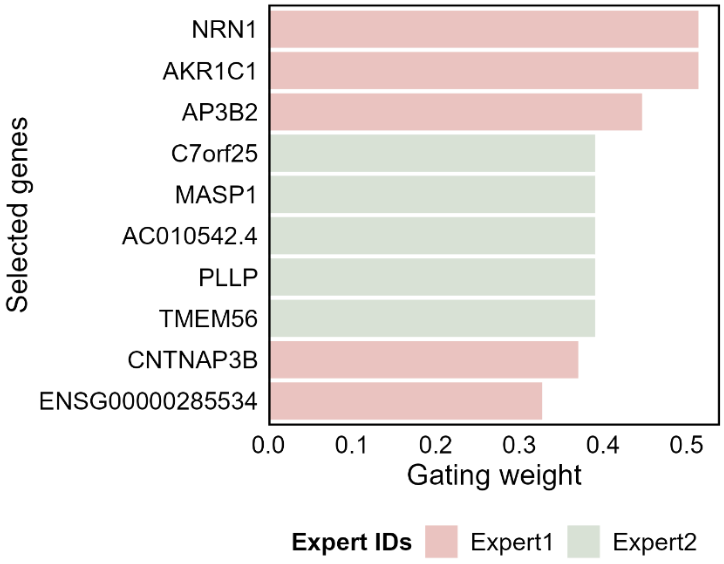


Supplementary Fig. 8 Top five prioritized genes per expert based on gating scores.

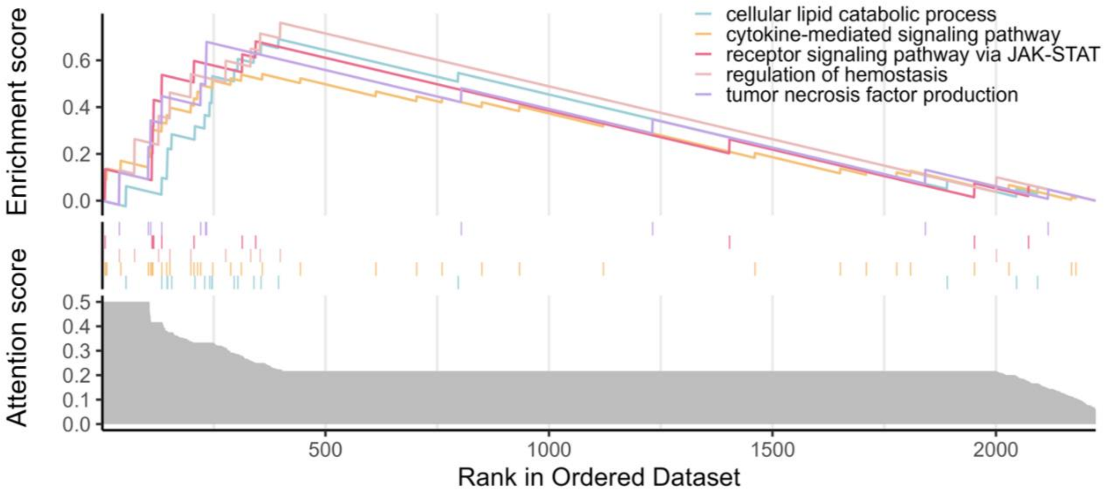


Supplementary Fig. 9 GSEA of genes ranked by expert-specific attention weights. Genes were rank-ordered according to attention weights derived from MEGA-ODE sub-experts. The GSEA results show enrichment of high-attention genes in representative COVID-19-related pathways.

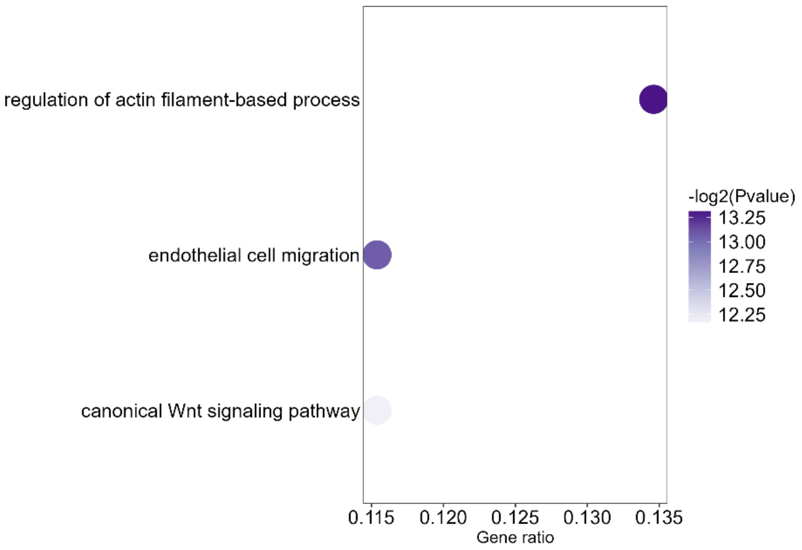


Supplementary Fig. 10 Gene Ontology enrichment analysis of top-ranked genes identified by gating scores.

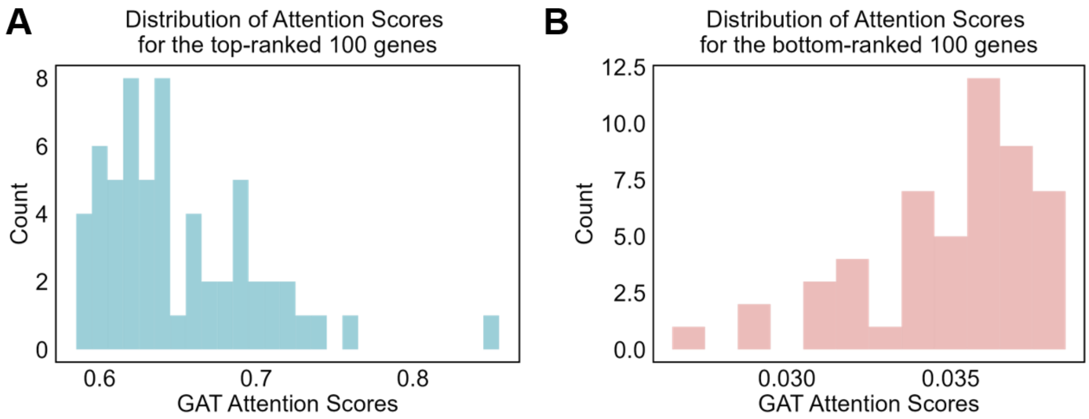


Supplementary Fig. 11 Distribution of attention scores for the top-ranked 100 and bottom-ranked 100 genes in the BMP4 perturbation dataset.

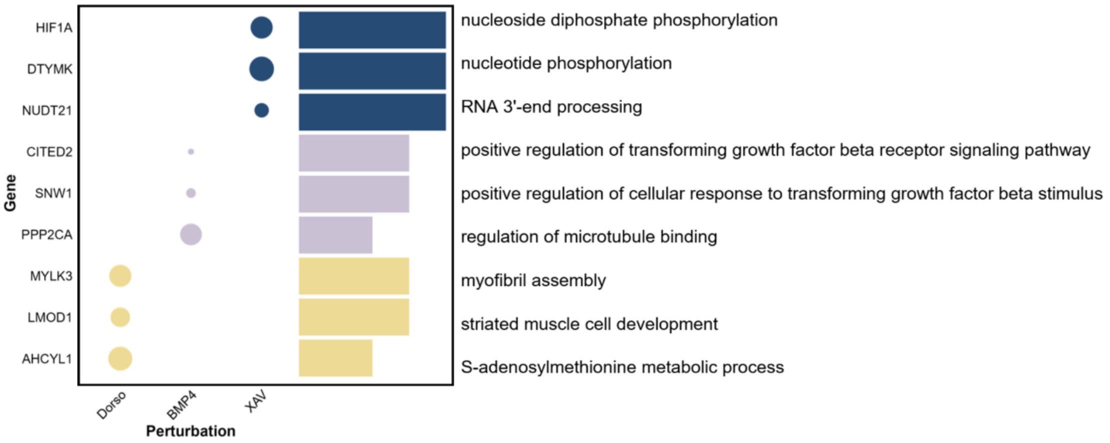


Supplementary Fig. 12 Integration of top-ranked genes and enriched GO terms across perturbations. The plot illustrates the relationship between prioritized genes and their functional roles under three perturbation conditions: Dorsomorphin, BMP4 and XAV. In the left bubble plot, the x-axis represents perturbation conditions, and the y-axis shows the top-ranked genes identified within each condition. Bubble size reflects the corresponding attention score. The bar plot on the right displays pathways enriched by Gene Ontology analysis of the corresponding genes.

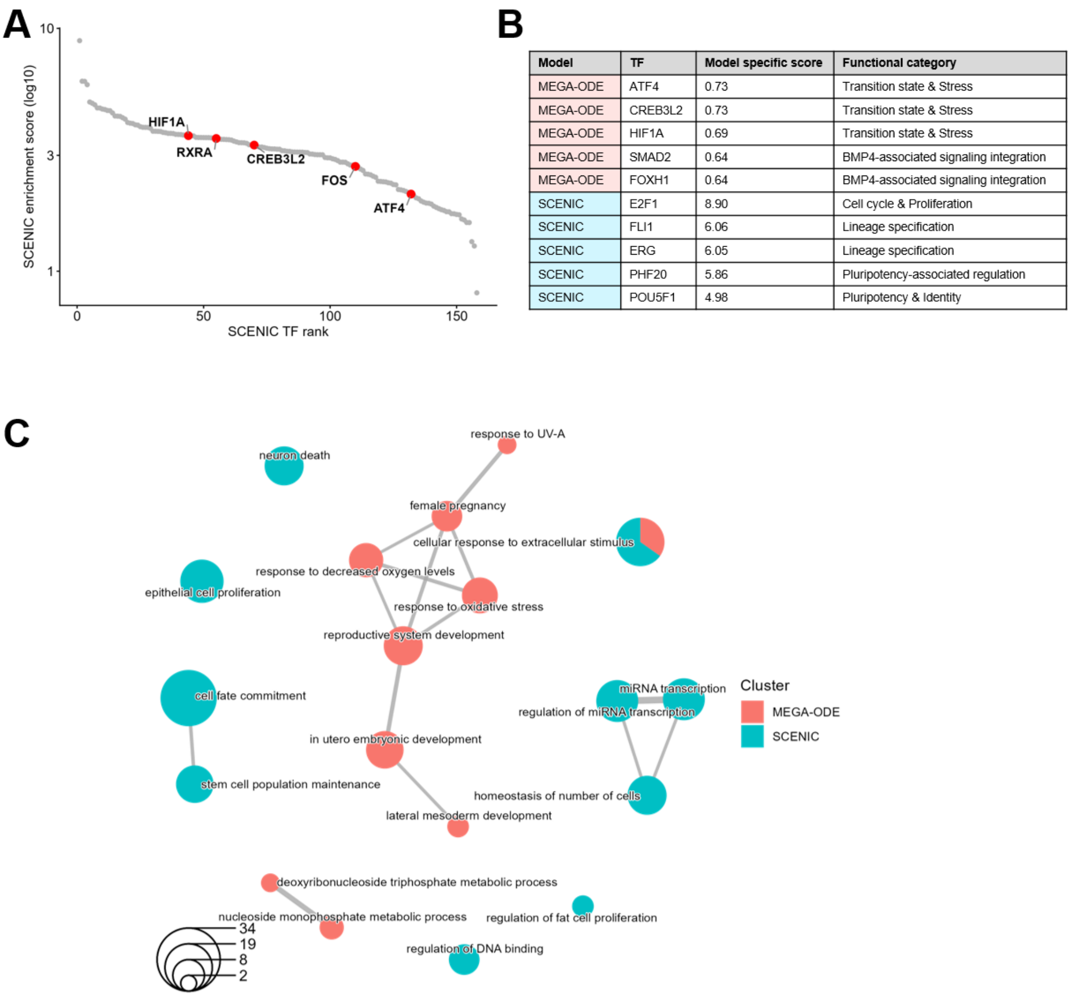


| Model | TF | Model specific score | Functional category |
|---|---|---|---|
| MEGA-ODE | ATF4 | 0.73 | Transition state & Stress |
| MEGA-ODE | CREB3L2 | 0.73 | Transition state & Stress |
| MEGA-ODE | HIF1A | 0.69 | Transition state & Stress |
| MEGA-ODE | SMAD2 | 0.64 | BMP4-associated signaling integration |
| MEGA-ODE | FOXH1 | 0.64 | BMP4-associated signaling integration |
| SCENIC | E2F1 | 8.90 | Cell cycle & Proliferation |
| SCENIC | FLI1 | 6.06 | Lineage specification |
| SCENIC | ERG | 6.05 | Lineage specification |
| SCENIC | PHF20 | 5.86 | Pluripotency-associated regulation |
| SCENIC | POU5F1 | 4.98 | Pluripotency & Identity |

Supplementary Fig. 13 Comparison of transcription factor prioritization between MEGA-ODE and SCENIC on BMP4-induced stem-cell differentiation data. (A) Overlapping TFs between MEGA-ODE and SCENIC. The x-axis shows the SCENIC TF rank, and the y-axis indicates SCENIC enrichment scores. Red points denote overlapping TFs between MEGA-ODE and SCENIC. (B) Summary of the top five TFs from each model, highlighting their scores and canonical roles. (C) Functional enrichment network. Biological process enrichment of genes from MEGA-ODE expert-specific top attention edges (110 genes) and SCENIC-inferred TFs (158 TFs), visualized as an enrichment map.

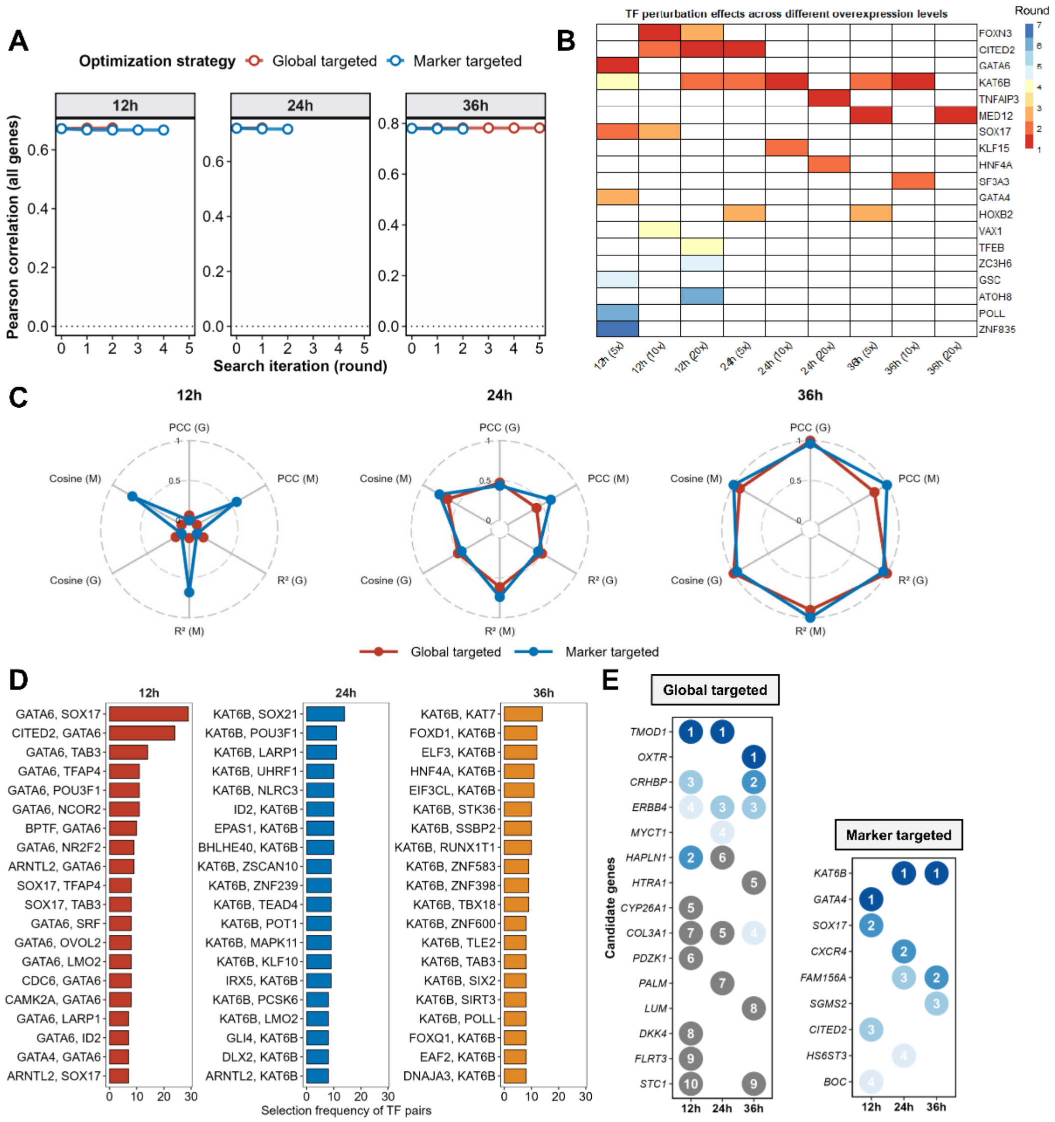


Supplementary Fig. 14 Extended analysis of in silico perturbation screening during hESC-to-definitive-endoderm differentiation. (A) Complementary analysis to the corresponding panel in Fig. 6C. The x-axis denotes the perturbation-screening time point across iterative search rounds, and the y-axis shows the Pearson correlation coefficient between the model-predicted expression of all 5,729 highly variable genes at 96 h and the observed 96 h expression profile. Colors indicate two objective functions, marker-targeted optimization and global-targeted optimization, enabling comparison of their search dynamics and predictive performance. (B) Candidate genes prioritized by marker-targeted optimization under different in silico overexpression magnitudes: 5×, 10× and 20×. This panel extends the main-text analysis by assessing the robustness of selected perturbation targets across perturbation strengths. (C) Distribution of multiple evaluation metrics under 10× overexpression for marker-targeted and global-targeted optimization. This analysis complements main-text Fig. 6C and D, where marker Pearson was shown as the primary readout. In the radar plots, metrics labeled with G were computed between the predicted and observed expression profiles across all highly variable genes, whereas metrics labeled with M were computed using only marker genes. (D) Extended TF-pair analysis corresponding to main-text Fig. 6F. Whereas the main figure shows only the top five TF pairs at each time point, this panel displays the top 20 TF pairs, providing a more comprehensive view of recurrent transcription factor combinations prioritized during perturbation screening.

(E) All-gene perturbation screening under 10× in silico overexpression. In contrast to the TF-restricted analyses in main-text Fig. 6B-F, this analysis expands the search space to all highly variable genes. The prioritized candidates include both transcription factors and non-TF genes.
